# Lack of Adipocyte Purinergic P2Y_6_ Receptor Greatly Improves Whole Body Glucose Homeostasis

**DOI:** 10.1101/2020.04.03.023234

**Authors:** Shanu Jain, Sai P. Pydi, Kiran S. Toti, Bernard Robaye, Marco Idzko, Oksana Gavrilova, Jürgen Wess, Kenneth A. Jacobson

**Affiliations:** Molecular Recognition Section, Laboratory of Bioorganic Chemistry, National Institute of Diabetes and Digestive and Kidney Diseases, Bethesda, MD 20892, USA; Molecular Signaling Section, Laboratory of Bioorganic Chemistry, National Institute of Diabetes and Digestive and Kidney Diseases, Bethesda, MD 20892, USA; Institute of Interdisciplinary Research, IRIBHM, Université Libre de Bruxelles, Gosselies, Belgium; Universitätsklinik für Innere Medizin II, Klinische Abteilung für Pulmologie, Medizinische Universität, Vienna, Austria and Dept. of Pneumology, and University Hospital Freiburg, Freiburg, Germany; Mouse Metabolism Core, National Institute of Diabetes and Digestive and Kidney Diseases, Bethesda, MD 20892, USA

## Abstract

Uridine diphosphate (UDP)-activated purinergic receptor P2Y_6_ (P2Y_6_R) plays a crucial role in controlling energy balance through central mechanisms. However, P2Y_6_R’s roles in peripheral tissues regulating energy and glucose homeostasis remain unexplored. Here, we report the surprising novel finding that adipocyte-specific deletion of P2Y_6_R protects mice from diet-induced obesity, improving glucose tolerance and insulin sensitivity with reduced systemic inflammation. These changes were associated with reduced JNK signaling, and enhanced expression and activity of PPARα affecting downstream PGC1α levels leading to beiging of white fat. In contrast, P2Y_6_R deletion in skeletal muscle reduced glucose uptake resulting in impaired glucose homeostasis. Interestingly, whole body P2Y_6_R KO mice showed metabolic improvements similar to those observed with mice lacking P2Y_6_R only in adipocytes. Our findings provide compelling evidence that P2Y_6_R antagonists may prove useful for the treatment of obesity and type 2 diabetes.

## INTRODUCTION

The increasing prevalence of obesity and type 2 diabetes (T2D) worldwide has led to added complications of cardiovascular disease, hypertension, fatty liver disease and cancer (*1, 2*). Disequilibrium between caloric intake and expenditure results in excess fat storage and hence the development of obesity (*3*). White adipose tissue (WAT) serves as the main site for the storage of excess calories as well as an active endocrine organ that regulates glucose and energy homeostasis in an organism (*4*). Conversely, brown adipose tissue (BAT) is a critical adipose depot responsible for non-shivering thermogenesis and energy expenditure (*4*). Adipose tissue undergoes extensive morphological and functional remodeling during obesity resulting in the development of insulin resistance, changing the repertoire of cytokines produced by the adipose tissues, and consequently increasing local and systemic inflammation (*5, 6*). Obesity-induced insulin resistance in other peripheral tissues such as skeletal muscle impairs glucose uptake causing hyperglycemia, thereby contributing to the core defect in T2D (*7*). Thus, there is an urgent need to define factors that affect the pathophysiology of adipose tissue and skeletal muscle controlling whole body (WB) energy balance and glycemic homeostasis. Identification of such factors may help to identify novel targets for the treatment of obesity and T2D.

Circulating nucleotides function as primary messengers in intercellular communication in an autocrine/paracrine manner (*8*). Nucleotides are released into the extracellular space in response to different pathophysiological conditions involving inflammation, cell lysis, hypoxia, trauma, and infection (*9, 10*). These agents play key roles in maintaining many important metabolic functions. High plasma levels of nucleoside/nucleotides such as adenosine and uridine have been reported under conditions of obesity, acting on G protein-coupled adenosine and P2Y receptors (P2YRs) of the purinergic receptor family (*11, 12*). P2Y receptors such as P2Y_1_R and P2Y_14_R have been shown to play a significant role in the regulation of leptin secretion and insulin sensitivity (*13, 14*). Activation of the G_q_-coupled P2Y_6_R by uridine diphosphate (UDP) increases glucose uptake in adipocyte 3T3-L1 and skeletal muscle C2C12 cell lines (*15*). P2Y_6_R also plays a pivotal role in inflammatory responses by stimulating immune cell migration and inducing the secretion of inflammatory cytokines such as MCP1 and IL6 (*16, 17*). Environmental stresses such as inflammation, chronic heart failure and dystrophic cardiomyopathy increase P2Y_6_R expression (*18, 19*). Expression of P2Y_6_R has also been reported in orexigenic agouti-related peptide expressing (AgRP)-neurons in the arcuate nucleus of the hypothalamus (*11*). P2Y_6_R-dependent activation of AgRP neurons promotes feeding in lean as well as obese mice (*11*). Blockade of P2Y_6_R reduced food intake and improved insulin sensitivity in obese mice (*20*).

In this study, we demonstrate that P2Y_6_R is expressed in adipose tissue and skeletal muscle and that its expression is regulated by obesity. We generated three different mouse models lacking P2Y_6_R throughout the body or selectively in mature adipocytes or skeletal muscle cells to decipher the receptor’s role in maintaining glucose and energy balance. Lack of P2Y_6_R specifically in adipocytes led to marked improvements in glucose and insulin tolerance in mice consuming an obesogenic diet. This improvement was notably due to impaired c-Jun N-terminal kinase (JNK) activation resulting in decreased systemic inflammation and increased browning in iWAT lacking P2Y_6_R.

Surprisingly, P2Y_6_R deletion specifically in skeletal muscle impaired glucose tolerance and insulin sensitivity. This impairment was due to decreased glucose uptake in skeletal muscle lacking P2Y_6_R. Lastly, global deletion of P2Y_6_R improved glucose metabolism and insulin tolerance along with reducing peripheral inflammation, suggesting that the lack of adipocyte P2Y_6_R is mainly responsible for this phenotype. Our data provide compelling evidence that attenuation of P2Y_6_R signaling in adipocytes can improve systemic glucose metabolism, suggesting that P2Y_6_R antagonists may be useful for the treatment of obesity and T2D.

## RESULTS

### P2Y6R is expressed in both adipose tissue and skeletal muscle, and is regulated differently in diet-induced obese mice

To investigate the potential role of P2Y_6_R in energy homeostasis and glucose metabolism, we quantified the mRNA expression levels of P2Y_6_R in metabolically important tissues such as adipose tissue and skeletal muscle. Of the three major adipose depots, P2Y_6_R mRNA expression was significantly higher in epididymal white adipose tissue (eWAT, visceral fat) than in inguinal white adipose tissue (iWAT, subcutaneous fat) and BAT **(Fig. 1A)**. Among the skeletal muscles examined, tibialis anterior muscle (TA) showed highest P2Y_6_R expression, followed by quadriceps (Q), gastrocnemius (G), and soleus (S) muscles **(Fig. 1B)**.

**Figure 1.**
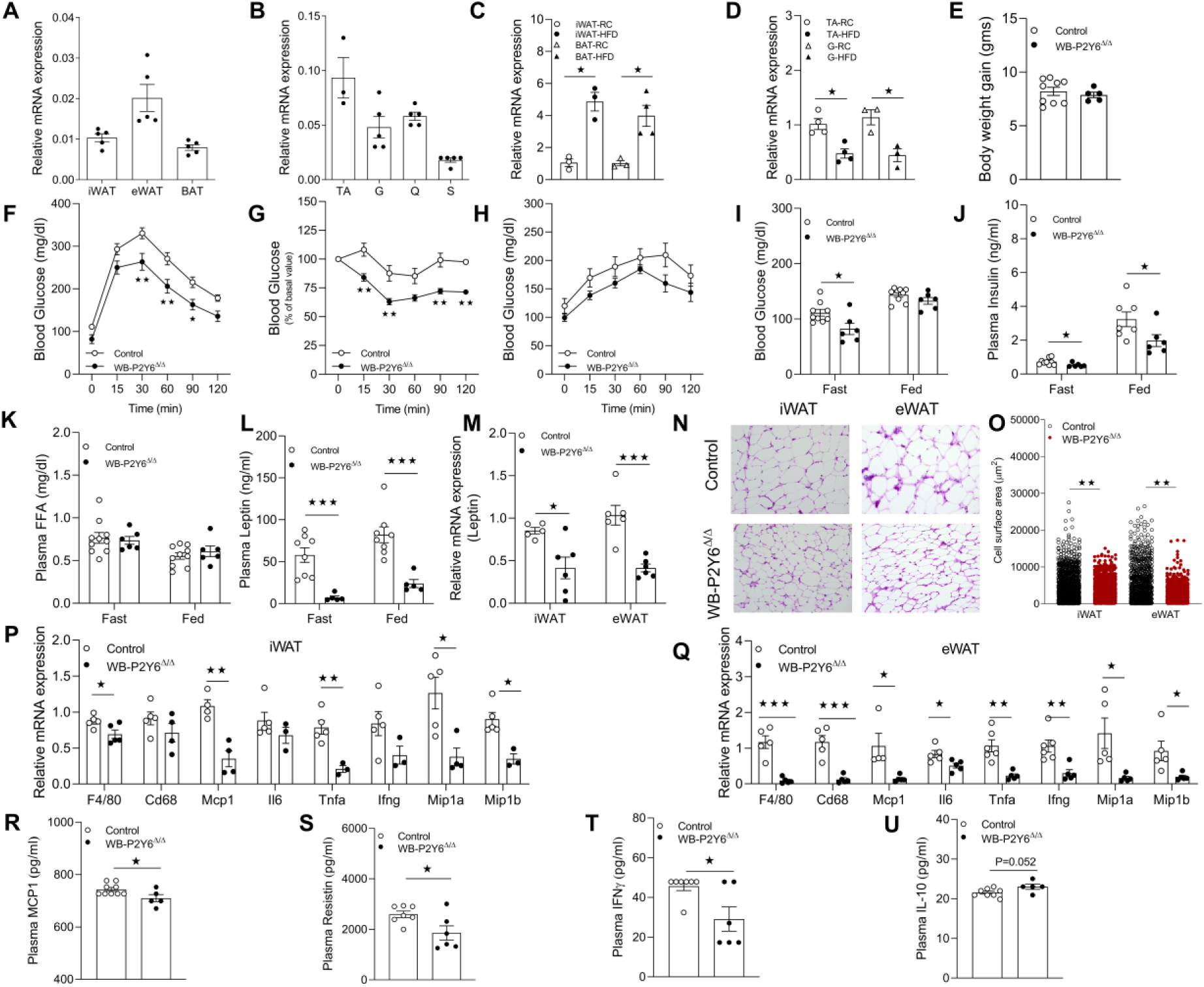
Whole body P2Y_6_R KO mice (WB-P2Y6^Δ/Δ^ mice) show reduced HFD-induced metabolic deficits and peripheral inflammation. (A) Relative expression of P2Y_6_R mRNA in subcutaneous (iWAT), visceral (eWAT) and brown adipose tissue (BAT) (n=5/group). (B) Relative expression of P2Y_6_R-mRNA in tibialis anterior (TA), gastrocnemius (G), quadriceps (Q) and soleus (S) skeletal muscle (n=3-5/group). (C) mRNA expression levels of P2Y_6_R in iWAT (mature adipocytes) or BAT from regular chow (RC) or high fat diet (HFD) fed C57BL/6 control mice. (n=3/group). (D) mRNA expression levels of P2Y_6_R in TA or G from regular chow (RC) or high fat diet (HFD) fed C57BL/6 control mice (n=3-4/group). (E) Body weight gain (grams) of mice maintained on HFD for 6 weeks (n=5-9/group). (F) Glucose tolerance test (IGTT, 1 g/kg glucose, i.p.) (n=6-9/group). (G) Insulin tolerance test (ITT, 1U/kg insulin, i.p.) (n=5-9/group). (H) Pyruvate tolerance test (PTT, 2 g/kg sodium pyruvate, i.p.) (n=5-9/group). (I) Fasting and fed blood glucose levels (n=6-9/group). (J) Fasting and fed plasma insulin levels (n=6-9/group). (K) Fasting plasma free fatty acid (FFA) levels (n=6-9/group). (L) Circulating levels of plasma leptin during fasting and fed state. (n=5-8/group). (M) mRNA levels of leptin in iWAT and eWAT of HFD fed WB-P2Y6^Δ/Δ^ and control mice (n=5-6/group). (N) Representative H&E-stained sections from iWAT and eWAT from WB-P2Y6^Δ/Δ^ and control mice on HFD. (O) Quantification data of adipocytes area in iWAT and eWAT from WB-P2Y6^Δ/Δ^ and control mice on HFD (n=10 sections/group). (P) mRNA expression levels of inflammation markers in iWAT of WB-P2Y6^Δ/Δ^ and control mice on HFD (n=3-6/group). (Q) mRNA expression levels of inflammation markers in eWAT of WB-P2Y6^Δ/Δ^ and control mice on HFD (n=3-6/group). (R-U) Circulating plasma levels of (R) MCP1, (S) resistin, (T) IFNγ, and (U) IL-10 in WB- P2Y6^Δ/Δ^ and control mice. (n=5-9/group). The expression of 18sRNA was used to normalize qRT-PCR data. All data are expressed as means ± SEM. *p< 0.05, **p< 0.01, ***p< 0.001 (C-E, I-M, O-U: two-tailed Student’s t-test; F-H: two-way ANOVA followed by Bonferroni’s post hoc test).

Obesogenic diets affect the expression of various G protein-coupled receptors (GPCRs) in distinct tissues (*21-23*). Therefore, we hypothesized that expression of P2Y_6_R may be altered in adipose tissue and skeletal muscle during obesity. To test the hypothesis, adipose tissues were collected from C57BL/6 mice fed regular chow or a high fat diet (HFD) for 8 weeks, and skeletal muscle tissues were collected from mice fed chow diet or HFD for 16 weeks. Interestingly, the HFD significantly increased P2Y_6_R mRNA expression in mature white adipocytes (free of immune cells) of iWAT and BAT **(Fig. 1C)**. To our surprise, P2Y_6_R expression was decreased in TA and G skeletal muscle of mice fed on HFD **(Fig. 1D)**, indicating a differential regulation of P2Y_6_R expression in the metabolic tissues under study.

### Global deletion of P2Y_6_R protects mice against HFD-induced metabolic deficits

As we observed marked differences in P2Y_6_R expression in metabolic tissues of HFD mice, we aimed to investigate the effect of whole-body deletion of P2Y_6_R (WB-P2Y6^Δ/Δ^) on the development of obesity and systemic glucose homeostasis. As expected, P2Y_6_R mRNA was undetectable in multiple metabolic tissues of WB-P2Y6^Δ/Δ^ mice, as compared to control mice (WB-P2Y6^WT/WT^) **(Fig. S1A)**. A group of control and WB-P2Y6^Δ/Δ^ mice (male C57BL/6 mice) was kept on HFD, and body weights were measured after 6 weeks on HFD. No significant differences in body weight gain was observed between groups **(Fig. 1E)**. Next, we subjected HFD WB-P2Y6^Δ/Δ^ and control mice to a series of metabolic tests. Interestingly, WB-P2Y6^Δ/Δ^ mice showed significant improvements in glucose tolerance and insulin sensitivity **(Fig. 1F, 1G)**. However, no differences in blood glucose levels were observed between the two groups in a pyruvate challenge test, indicating that hepatic glucose output was similar in the two groups **(Fig. 1H)**. WB-P2Y6^Δ/Δ^ mice exhibited reduced fasting blood glucose levels **(Fig. 1I)** and reduced plasma insulin levels under both fasting and fed conditions (**Fig. 1J)**, indicating improved peripheral insulin sensitivity. Plasma FFA levels did not differ between the two mouse cohorts, indicating that P2Y_6_R does not play a significant role in adipose tissue lipolysis **(Fig. 1K)**. Interestingly, we observed significant decrease in the plasma leptin levels in in WB-P2Y6^Δ/Δ^ mice under both fasting and fed conditions **(Fig. 1L)**. In agreement with reduced plasma leptin levels, leptin mRNA levels were significantly decreased in in iWAT and eWAT of WB-P2Y6^Δ/Δ^ mice **(Fig. 1M)**. Deficiency of P2Y_6_R did not cause any significant changes in the expression of genes involved in adipocyte browning/mitochondrial function in iWAT and eWAT of WB- P2Y6^Δ/Δ^, as compared to tissues from control littermates **(Fig. S1B, 1C)**. Surprisingly, BAT lacking P2Y_6_R showed modest decreases in mRNA levels of genes such as *Ucp1, Pparα* and *Pparg* **(Fig. S1D)**.

### Reduced inflammation in global P2Y_6_R-deficient mice

Development of obesity and subsequent insulin resistance is associated with enhanced systemic inflammation (*5, 24*). Activation of P2Y_6_R by UDP has been shown to mediate inflammatory responses by increasing the secretion of pro-inflammatory cytokines such as monocyte chemoattractant protein-1 (MCP1) (*16*). Therefore, we hypothesized that global deletion of P2Y_6_R may protect mice from peripheral inflammation resulting from HFD consumption. To test this hypothesis, H&E staining was performed on iWAT and eWAT sections of HFD WB- P2Y6^Δ/Δ^ and control mice. eWAT sections from control mice with a functional P2Y_6_R exhibited crown-like structures indicating infiltration of inflammatory monocyte/macrophages. Such structures were reduced in eWAT sections of WB-P2Y6^Δ/Δ^ mice (**Fig. 1N)**. Imaging studies also revealed an increased number of smaller adipocytes in both iWAT and eWAT of WB-P2Y6^Δ/Δ^ compared to control tissues **(Fig. 1N, 1O)**. A similar phenomenon was observed with BAT sections **(Fig. S1D)**. To further confirm these findings, mRNA levels of pro-inflammatory markers were quantified in different adipose depots. As expected, mRNA levels of macrophage markers (*F4/80* and *Cd68*) were significantly reduced in iWAT and eWAT of WB-P2Y6^Δ/Δ^ mice **(Fig. 1P,1 Q)**. Transcript levels of pro-inflammatory markers such as *Mcp1, Il6, Tnfα, Ifng, Mip1a* and *Mip1b* were also significantly reduced in adipose depots of WB-P2Y6^Δ/Δ^ mice **(Fig. 1P, 1Q, Fig S1F-1H)**. In accordance with the gene expression data, circulating plasma levels of MCP1, resistin and IFNγ were significantly lower in WB-P2Y6^Δ/Δ^ mice compared to control littermates **(Fig. 1R-1U)**. Taken together, this data clearly indicates that global deletion of P2Y_6_R reduced peripheral inflammation thereby improving glucose homeostasis and insulin sensitivity.

### Pharmacological determination of functional P2Y_6_R expression in mouse white and brown adipocytes

We previously identified uracil nucleotide derivatives that act as potent human P2Y_6_R (hP2Y_6_R) agonists (*25*). We selected the potent P2Y_6_R agonist MRS4383 (((1*S*,2*R*,3*R*,4*S*,5*R*)-3,4- dihydroxy-5-((*Z*)-4-(methoxyimino)-2-oxo-3,4-dihydropyrimidin-1(2*H*)-yl)bicyclo[3.1.0]hexan- 2-yl)methyl trihydrogen diphosphate) to identify and demonstrate a functional P2Y_6_R expressed by mouse (m) adipocytes. Specifically, we first isolated the stromal vascular fraction (SVF) from both iWAT and BAT of WB-P2Y6^Δ/Δ^ and control mice and then differentiated the cells into mature white or brown adipocytes. Treatment of mature adipocytes with different concentrations of MRS4383 resulted in accumulation of inositol monophosphate (IP1) in control cells expressing P2Y_6_R **(Fig. 2A, 2B)**. In contrast, this response was absent in cells lacking functional P2Y_6_R **(Fig. 2A, 2B)**. This experiment confirmed the specificity of MRS4383 for the G_q_-coupled P2Y_6_R and the presence of functional P2Y_6_Rs in mouse white and brown adipocytes.

**Figure 2.**
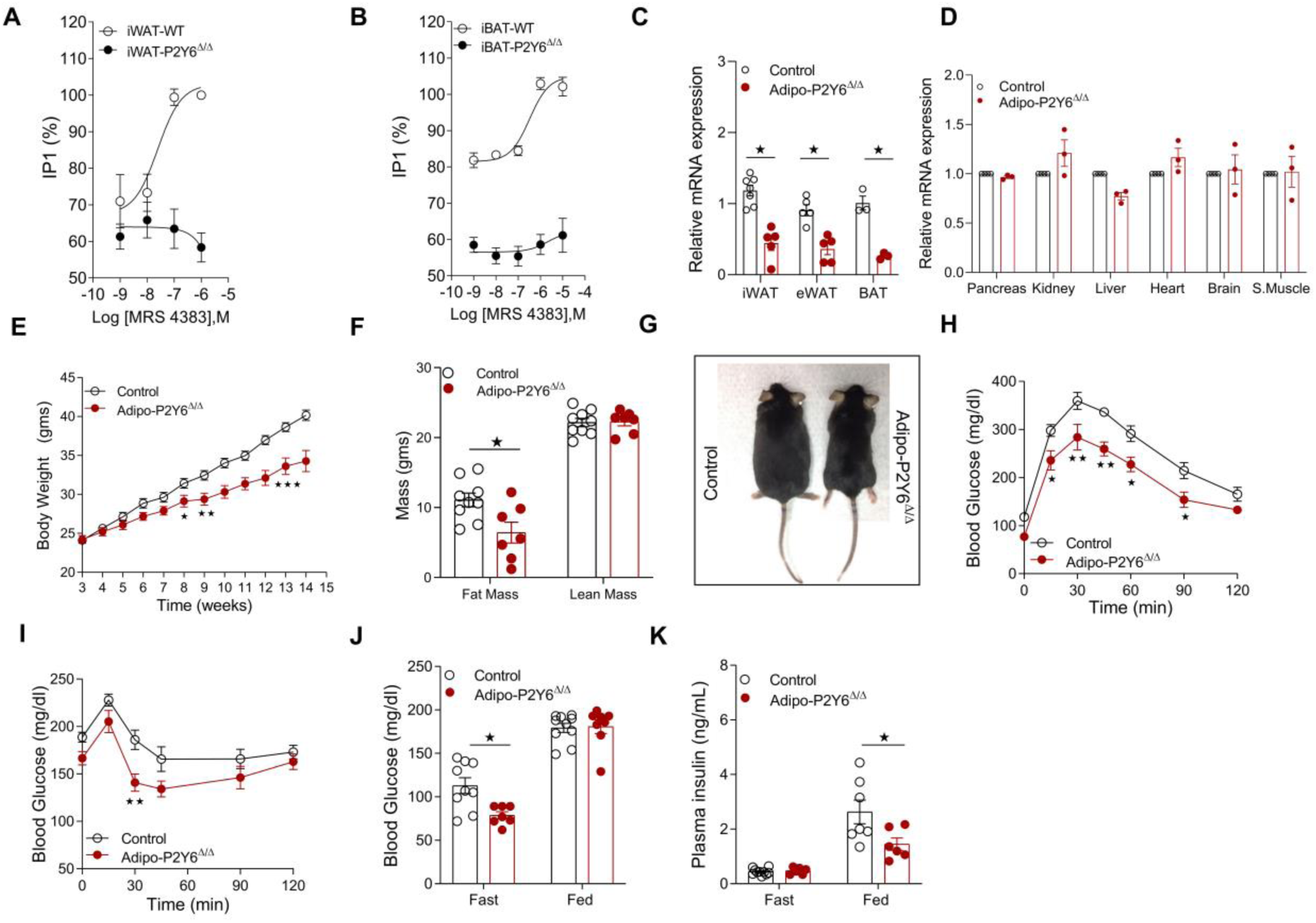
Adipocyte-specific P2Y_6_R KO mice (adipo-P2Y6^Δ/Δ^ mice) are protected from HFD-induced obesity and metabolic deficits. (A) IP1 accumulation assay in the differentiated mature white adipocytes from WB-P2Y6^Δ/Δ^ and control mice. (n=3-4), each experiment was performed in triplicate (B) IP1 accumulation assay in differentiated mature brown adipocytes from WB-P2Y6^Δ/Δ^ and control mice (n=3-4). Each experiment was performed in triplicate. (C) mRNA levels of P2Y_6_R in mature adipocytes isolated from iWAT (n=5-7/group), eWAT (n=5/group) and BAT (n=3/group) of adipo-P2Y6^Δ/Δ^ and control mice. (D) mRNA levels of P2Y_6_R in pancreas, kidney, liver, heart, brain and skeletal muscle of adipo-P2Y6^Δ/Δ^ and control mice (n=3/group). (E) Body weight measurements of mice maintained on HFD (n=7-9/group). (F) Body composition (lean and fat mass in g) of mice maintained on HFD (n=7-9/group). (G) Images of representative adipo-P2Y6^Δ/Δ^ and control mice (10 weeks on HFD). (H) Glucose tolerance test (IGTT, 1g /kg glucose, i.p.) (n=6-8/group). (I) Insulin tolerance test (ITT, 1U/kg insulin, i.p (n=6-8/group). (J) Fasting and fed blood glucose levels (n=7-10/group). (K) Fasting plasma insulin levels (n=6-8/group). The expression of 18sRNA was used to normalize qRT-PCR data. All data are expressed as means ± SEM. *p< 0.05, **p< 0.01, ***p< 0.001 (C-F, J-K: two-tailed Student’s t-test; H-I: two-way ANOVA followed by Bonferroni’s post hoc test). All experiments were conducted on mice fed on HFD.

### Generation of adipocyte-specific P2Y_6_R KO mice (adipo-P2Y6^Δ/Δ^)

Given that global deletion of P2Y_6_R improves glucose metabolism with reduced peripheral inflammation, we aimed to investigate whether abrogation of P2Y_6_R signaling in mature adipocytes affects energy and glucose homeostasis. To this end, we inactivated the P2Y_6_R gene selectively in mature adipocytes by crossing floxed P2Y_6_R (*P2Y6f/f*) mice with mice expressing Cre recombinase under the control of the adipocyte-specific adiponectin promoter (*adipoq-Cre* mice). This mating scheme gave rise to *P2Y6f/f* (control) and *adipoq-Cre*^*+/-*^*P2Y6f/f* (adipo- P2Y6^Δ/Δ^) mice. Real-time qPCR analysis revealed a significant knockdown of P2Y_6_R mRNA in mature adipocytes from different adipose depots (iWAT, eWAT, BAT) of adipo-P2Y6^Δ/Δ^ mice **(Fig. 2C)**. P2Y_6_R expression remained unaltered in other metabolically active tissues **(Fig. 2D)**.

### Adipo-P2Y6^Δ/Δ^ mice are protected from HFD-induced obesity and metabolic dysfunction

To assess the metabolic roles of adipocyte P2Y_6_R, adipo-P2Y6^Δ/Δ^ mice and control littermates were first maintained on regular chow diet. Under these conditions, the two groups of mice showed similar body weight, glucose tolerance, insulin sensitivity, blood glucose, plasma glycerol, FFA and triglyceride levels in either the fasting or fed state **(Fig. S2A-2G)**. However, under fasting conditions, plasma insulin levels were reduced in adipo-P2Y6^Δ/Δ^ mice, while no difference was observed in the fed state **(Fig. S2H)**. Similarly, plasma leptin levels were reduced in adipo-P2Y6^Δ/Δ^ mice **(Fig. S2I)**.

Next, to determine the potential role of adipocyte P2Y_6_R in diet-induced dysregulation of energy homeostasis and glucose metabolism, adipo-P2Y6^Δ/Δ^ and control mice were challenged with a HFD. Adipo-P2Y6^Δ/Δ^ mice gained less body weight than control mice **(Fig. 2E)**, and MRI scanning data revealed that differences in body weight were due to a significant reduction in fat mass in adipo-P2Y6^Δ/Δ^ mice **(Fig. 2F, 2G)**. Next, a series of metabolic tests were carried out with both groups of mice maintained on HFD for 8 weeks. Under these conditions, adipo- P2Y6^Δ/Δ^ mice showed significant improvements in glucose tolerance **(Fig. 2H)** and insulin sensitivity **(Fig. 2I)**. Moreover, blood glucose levels (fasting) and plasma insulin levels (fed) were significantly reduced in adipo-P2Y6^Δ/Δ^ mice **(Fig. 2J, 2K)**. Loss of adipocyte P2Y_6_R had no significant effect on adipocyte lipolysis as plasma FFA, glycerol, and total triglyceride levels were similar between the two groups. **(Fig. S3A-3C)**.

To examine the effect of lack of P2Y_6_R in adipocytes on whole-body energy homeostasis, we performed indirect calorimetry measurements **(Fig. S3)**. The measurements were performed on mice during the first week (first 4 days) of HFD feeding. Adipo-P2Y6^Δ/Δ^ mice showed no significant differences in total energy expenditure (TEE), food intake, oxygen consumption rate, and respiratory exchange ratio **(Fig S3D-3G)**.

### Adipo-P2Y6^Δ/Δ^ mice showed reduced inflammation on HFD

As global deletion of P2Y_6_R ameliorates inflammation associated with HFD feeding, we investigated whether deletion of P2Y_6_R specifically in mature adipocytes affected peripheral inflammation. To test this hypothesis, H&E stained iWAT, eWAT and BAT sections from HFD adipo-P2Y6^Δ/Δ^ and control mice were analyzed microscopically (10X magnification). Adipose tissue (eWAT) from control mice showed pronounced infiltration of pro-inflammatory immune cells **(Fig. 3A)**. This effect was significantly reduced in eWAT of HFD adipo-P2Y6^Δ/Δ^ mice **(Fig. 3A)**. Consistent with the results obtained with WB-P2Y6^Δ/Δ^ mice, adipo-P2Y6^Δ/Δ^ mice also showed significantly smaller adipocyte size, as compared to the corresponding adipose tissues from control littermates **(Fig. 3A, 3B)**.

**Figure 3:**
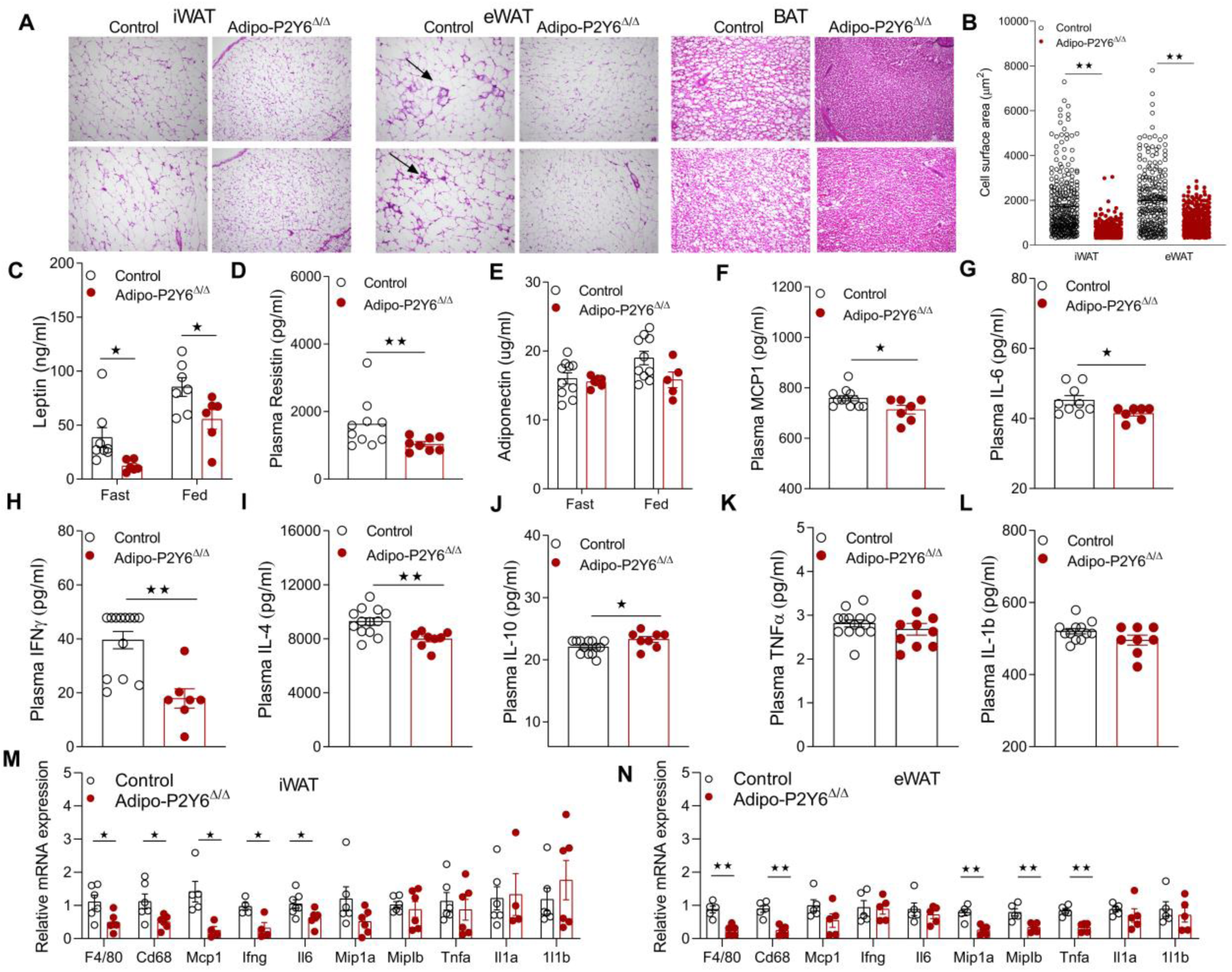
Reduced inflammation in adipo-P2Y6^Δ/Δ^ mice. (A) Representative H&E-stained sections from iWAT, eWAT and BAT from adipo-P2Y6^Δ/Δ^ and control mice maintained on HFD. (B) Cell surface area quantified using Adiposoft software in iWAT and eWAT sections. (n=5 sections/group). (C-L) Circulating plasma levels of (C) leptin, (D) resistin, (E) adiponectin, (F) MCP1, (G) IL6, (H) IFNγ, (I) IL4, (J) IL10, (K) TNFα, and (L) IL1b in adipo-P2Y6^Δ/Δ^ and control mice (n=6-12/group). (M, N) mRNA expression levels of inflammatory markers in (M) iWAT and (N) eWAT from adipo-P2Y6^Δ/Δ^ and control mice (n=4-6/group). The expression of 18sRNA was used to normalize qRT-PCR data. All data are expressed as means ± SEM. *p< 0.05, **p< 0.01, ***p< 0.001 (B-N: two-tailed Student’s t-test). All experiments were conducted on mice fed on HFD.

Under conditions of obesity and T2D, the secretion of adipokines/cytokines from adipocytes changes drastically, resulting in significant effects on whole body glucose and energy metabolism (*6, 26*). P2Y_6_R has been shown to regulate the secretion of cytokines mediating inflammation under pathophysiological conditions (*27*). Therefore, we measured circulating levels of adipokines/cytokines in adipo-P2Y6^Δ/Δ^ mice. Plasma leptin and resistin levels were reduced in adipo-P2Y6^Δ/Δ^ mice, consistent with reduced obesity (**Fig. 3C, 3D**). No difference in adiponectin levels was found between the groups **(Fig. 3E)**. Among the pro-inflammatory cytokines, plasma levels of MCP1, IL6, IFNγ and IL4 were significantly decreased, while the levels of IL10, an anti-inflammatory cytokine, were increased in adipo-P2Y6^Δ/Δ^ mice **(Fig 3F-3L)**. Consistent with staining data, mRNA expression levels of macrophage markers (*F4/80* and *Cd68*) were significantly reduced in iWAT, eWAT and BAT from adipo-P2Y6^Δ/Δ^ mice **(Fig. 3M, 3N and Fig. S3H)**. In agreement with the plasma cytokine levels, iWAT showed decreased transcript levels of *Mcp1, Ifng* and *Il6*, while eWAT displayed reduced gene expression of *Mip1a, Mip1b* and *Tnfa* **(Fig. 3M-3N)**.

Collectively, these experiments revealed that either global deletion or adipocyte-specific deficiency of P2Y_6_R protects against obesity-associated inflammation and improves glucose and insulin tolerance in mice.

### Reduced hepatic steatosis and hepatic inflammation in adipo-P2Y6^Δ/Δ^ mice

Since adipo-P2Y6^Δ/Δ^ mice showed reduced adiposity but no change in plasma FFA levels, we aimed to understand the effect of reduced fat mass on the ectopic deposition of fat in tissues like liver. Liver sections from HFD adipo-P2Y6^Δ/Δ^ and control mice were subjected to H&E and Oil red O staining. Microscopic analysis revealed reduced deposition of fat in the liver of adipo- P2Y6^Δ/Δ^ mice **(Fig S4A)**. Consistent with the observation, liver weight as well as liver triglyceride levels were decreased in the adipo-P2Y6^Δ/Δ^ mice **(Fig. S4B and S4C)**. The decrease in fat deposition resulted in reduced hepatic inflammation as evident from the reduced mRNA levels of inflammatory markers (*Il6, Ifng, Mcp1, Mip1b and Il1b*) in the liver of adipo-P2Y6^Δ/Δ^ mice, contributing to reduced peripheral inflammation **(Fig. S4D)**. However, no significant differences in expression levels of genes involved in the hepatic glucose and lipid metabolism were observed **(Fig S4E)**.

### P2Y_6_R-mediated JNK activation in mature white adipocytes

G_q_ protein-dependent activation of JNK signaling has been reported previously (*28*). JNK activation in adipocytes has been shown to result in metabolic dysfunction (*29*). Hence, we hypothesized that P2Y_6_R-mediated signaling may affect JNK activation in adipocytes and contribute to the observed metabolic phenotype. To this end, preadipocytes from iWAT of WB- P2Y6^Δ/Δ^ and control mice were isolated and differentiated into mature adipocytes. Mature adipocytes were stimulated with the P2Y_6_R-selective agonist MRS4383 for specific periods of time, and JNK phosphorylation was quantified. Strikingly, P2Y_6_R activation caused phosphorylation of JNK in control adipocytes but not in WB-P2Y6^Δ/Δ^ derived adipocytes **(Fig. 4A)**. Adipocytes lacking P2Y_6_R also showed decreased levels of p-cJUN, a downstream effector of JNK **(Fig. 4B)**. We also quantitated protein levels of total JNK (T-JNK) and p-JNK in iWAT from adipo-P2Y6^Δ/Δ^ and control mice. Strikingly, iWAT from adipo-P2Y6^Δ/Δ^ mice showed decreased levels of T-JNK and p-JNK compared to control mice **(Fig. 4C, 4D)**, confirming our hypothesis that activation of P2Y_6_R regulates JNK and cJUN.

**Figure 4:**
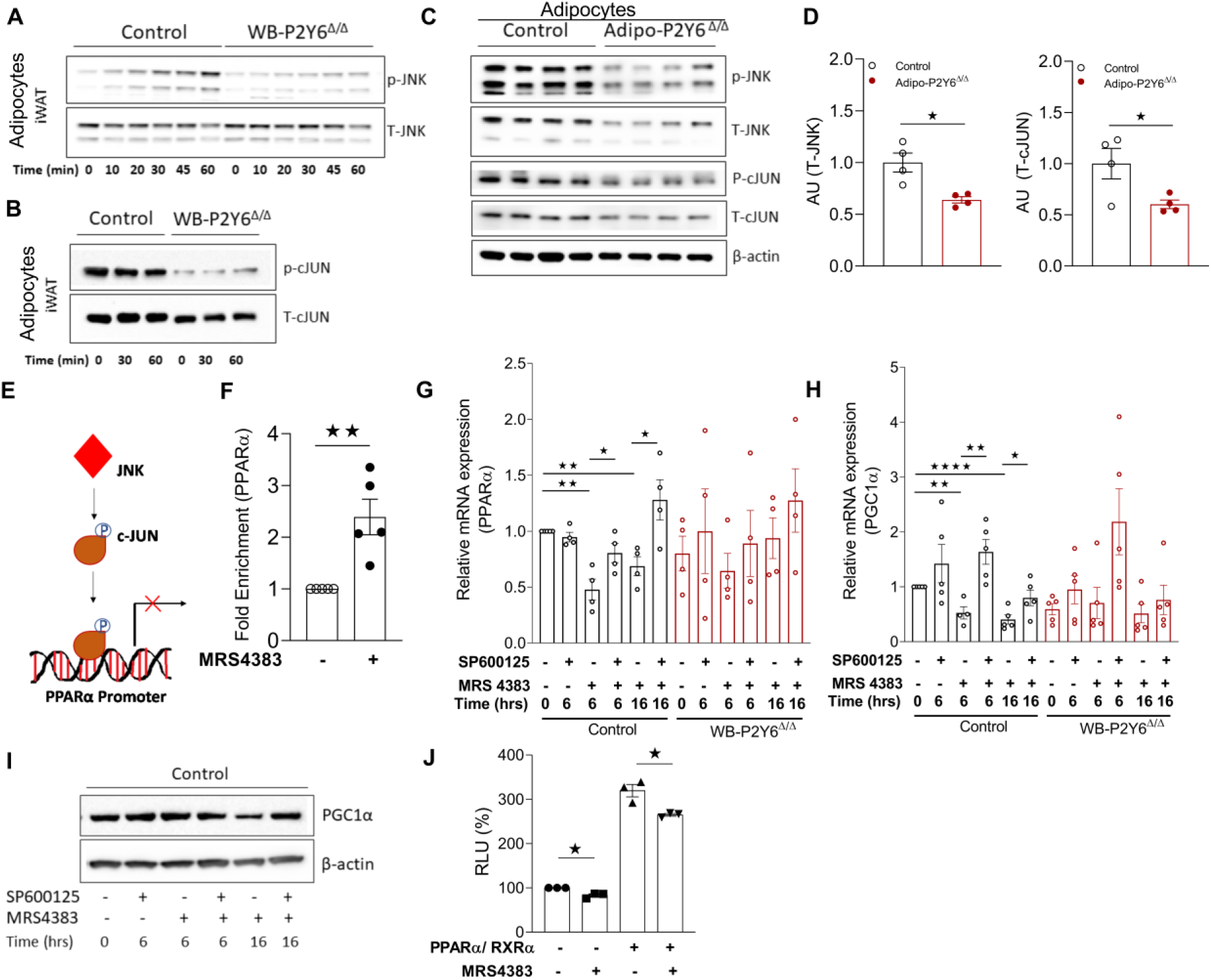
P2Y_6_R regulates JNK-PPARα-PGC1α axis in white adipocytes. (A) Western blot analysis of p-JNK/T-JNK protein levels in mature adipocytes differentiated from iWAT preadipocytes isolated from WB-P2Y6^Δ/Δ^ and control mice. Mature adipocytes were treated with P2Y_6_R agonist, MRS4383 (1 µM), and cells were collected at the indicated time points. Representative blots are shown (n=4 independent experiments). (B) p-cJUN/T-cJUN protein expression was measured using western blot analysis in iWAT mature adipocytes differentiated *in-vitro* using preadipocytes isolated from WB-P2Y6^Δ/Δ^ and control mice. Mature adipocytes were treated with P2Y_6_R agonist, MRS4383 (1 µM), and cells were collected at indicated time points. Representative blot are shown (n=3 independent experiments). (C) Western blots showing increased T-JNK, p-JNK, T-cJUN and p-cJUN expression in iWAT of HFD fed adipo-P2Y6^Δ/Δ^ and control mice. Each lane represents a different mouse (n=4/group). (D) Quantification of immunoblotting data shown in panel (C). (E) Schematic representation of the mechanism by which P2Y_6_R activation regulates the interaction between c-JUN and the *PPARα* promoter. (F) Chip assay showing fold enrichment of *PPARα* promoter region in mature adipocytes differentiated from iWAT preadipocytes isolated from WB-P2Y6^Δ/Δ^ and control mice. Mature adipocytes were treated with P2Y_6_R agonist, MRS4383 (1 µM) and cell were collected at after 2_1/2 h_. (n=5 independent experiments). (G, H) mRNA expression analysis of *Ppara* and *Pgc1a* genes in differentiated iWAT adipocytes isolated from WB-P2Y6^Δ/Δ^ and control mice. Mature adipocytes were incubated with JNK inhibitor (SP600126, 25 µM) and/or P2Y_6_R agonist (MRS4383, 1 µM) for the indicated periods of time. Cells were collected at specific time points as indicated (n=4/group). (I) Western blot analysis for PGC1α protein expression in differentiated iWAT adipocytes isolated from WB-P2Y6^Δ/Δ^ and control mice. A representative blot is shown (n=3 independent experiments). (J) PPARα activity assay using a luciferase assay in 1321N1 astrocytoma cells expressing h P2Y_6_R. Transfected cells were treated with MRS4383, 1 µM for 6 h. Relative luciferase activity was normalized to β-galactosidase activity for each sample (n=3 independent experiments). The expression of 18sRNA was used to normalize qRT-PCR data. All data are expressed as means ± SEM. *p< 0.05, **p< 0.01, ***p< 0.001 (two-tailed Student’s t-test). RLU, Relative luminescence units. All preadipocytes were isolated from RC fed mice. Preadipocytes were differentiated to mature adipocytes for experiments.

### P2Y_6_R regulates the JNK-PPARα-PGC1α axis in white adipocytes

To further delineate the contribution of P2Y_6_R mediated JNK signaling in the regulation of adipocyte metabolism, we identified a signaling cascade downstream of P2Y_6_R in adipocytes. In cardiac myocytes, transcription of PPARα is regulated by binding of cJUN to the PPARα promoter region (*30*). Since P2Y_6_R activation regulates JNK, we investigated whether JNK-mediated activation of cJUN affected the expression of PPARα. To this end, we performed a chromatin immunoprecipitation (ChIP) assay using nuclear DNA from adipocytes expressing P2Y_6_R with or without stimulation with P2Y_6_R agonist (MRS4383, 1 µM). P2Y_6_R activation led to an increase in the enrichment of PPARα promoter sequence in DNA isolated from cJUN antibody-bound chromatin regions **(Fig. 4E, 4F)**. To further demonstrate that P2Y_6_R mediated JNK activation can regulate PPARα, we treated cultured mature adipocytes from WB-P2Y6^Δ/Δ^ and control mice with JNK inhibitor (SP600125, 25 µM) and then stimulated with P2Y_6_R agonist (MRS4383, 1 µM) for the indicated time period. Treatment of cells with the JNK inhibitor alone had no effect on *Ppara* mRNA levels in adipocytes **(Fig 4G)**. However, treatment of cells with P2Y_6_R agonist decreased *Ppara* mRNA levels in control cells but not in adipocytes lacking P2Y_6_R expression, indicating P2Y_6_R specific regulation of *Ppara* levels **(Fig 4G)**. The observed decrease in *Ppara* levels by P2Y_6_R activation could be prevented by pretreatment of cells with JNK inhibitor **(Fig. 4G)**. P2Y_6_R activation in adipocytes also decreased *PGC1α* mRNA levels, a coactivator of PPARα in the transcriptional control of mitochondrial genes **(Fig. 4H)**. Consistent with the change in mRNA levels, the level of PGC1α protein was significantly decreased after 16 h of P2Y_6_R activation. Prior treatment of adipocytes with JNK inhibitor prevented the decrease in PGC1α protein levels **(Fig. 4I)**. Taken together, our data indicate that JNK-mediated cJUN activation promotes the cJUN interaction with the PPARα promoter, regulating the expression of PPARα and its downstream target and co-activator such as PGC1α.

Further, we examined PPARα activity following P2Y_6_R activation using a luciferase reporter assay. PPARα forms a heterodimer with the retinoid X receptor (RXR) that binds to their DNA binding element known as PPAR response element (PPRE)/Direct repeat 1 (DR1) (*31*). Astrocytoma 1321N1 cells expressing hP2Y_6_R were transiently transfected with DR1 plasmid with or without plasmids expressing PPARα/RXR. Stimulation of transfected cells with MRS4383 (1 µM) decreased the luciferase activity, indicating that decreased PPARα activity was due to activation of P2Y_6_R signaling in these cells **(Fig. 4J)**.

### Ablation of P2Y_6_R in iWAT promotes the expression of mitochondrial/ beiging specific genes

To recapitulate the effects of P2Y_6_R signaling observed in primary adipocytes, whole-transcriptome analysis of iWAT from HFD adipo-P2Y6^Δ/Δ^ and control mice was performed. Analysis of differentially expressed genes revealed that ∼ 406 genes were up-regulated whereas ∼ 470 genes were downregulated in iWAT of adipo-P2Y6^Δ/Δ^ mice (Genomatix Genome Analyzer, cut off: log2 (fold change) ≥ 0.58, p-value 0.05) **(Fig. 5A, Table S1)**. Gq-protein mediated pathways such as cellular calcium fluxes and actin-cytoskeletal signaling were found regulated in the iWAT of adipo-P2Y6^Δ/Δ^ mice **(Fig. 5B)**.

**Figure 5:**
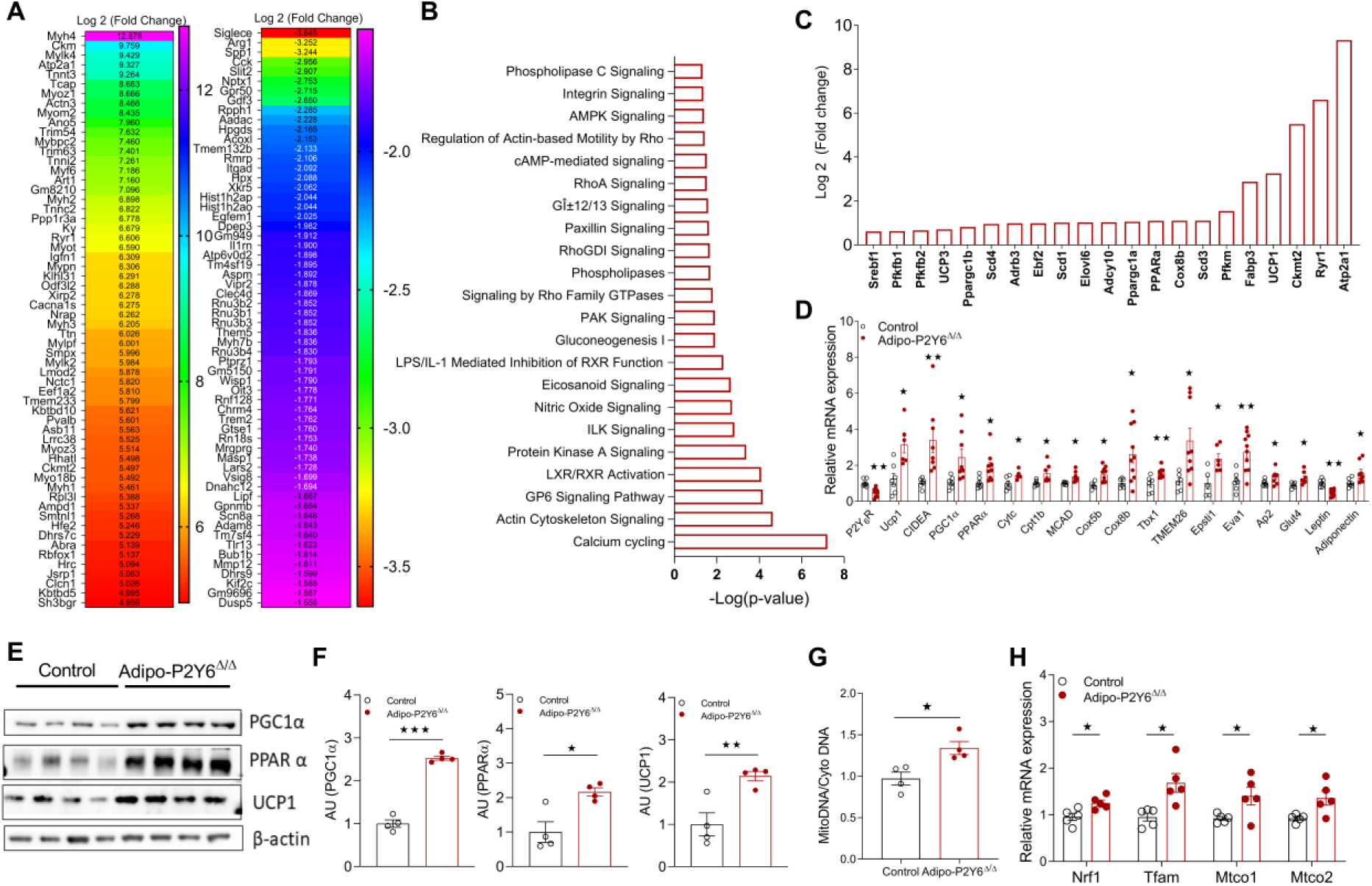
iWAT remodeling to beige phenotype in adipo-P2Y6^Δ/Δ^ mice. (A) RNA-seq analysis showing the 60 top upregulated and 60 top downregulated genes in iWAT of HFD adipo-P2Y6^Δ/Δ^ compared to HFD control littermates. Data are represented as a heat map of genes where color gradation corresponds to the log 2 (fold change) (n=6 libraries/group). (B) Column representation of significantly regulated pathways for all differentially expressed genes in iWAT of adipo-P2Y6^Δ/Δ^ mice (p<0.05, Ingenuity pathway analysis (IPA)) (n=6 libraries/group). (C) Log2 (Fold change) of genes involved in imparting beige-like phenotype to iWAT of adipo-P2Y6^Δ/Δ^ mice (p<0.05) (n=6 libraries/group). (D) RT-PCR for verifying changes in mRNA expression of browning/mitochondrial markers in iWAT of adipo-P2Y6^Δ/Δ^ mice compared to the control group (n=4-6 per group). (E) Western botting analysis of PGC1α, PPARα and UCP1 protein levels in iWAT of adipo- P2Y6^Δ/Δ^ and control mice (n=4/group). (F) Quantification of immunoblotting data shown in panel (E) (n=4/group). (G) MitoDNA (mitochondrial DNA)/CytoDNA (cytoplasmic DNA) in iWAT of adipo- P2Y6^Δ/Δ^ and control mice (n=4/group). (H) mRNA expression levels of mitochondrial biogenesis markers in the iWAT of adipo- P2Y6^Δ/Δ^ and control mice (n=4-6/group). The expression of 18sRNA was used to normalize qRT-PCR data. All data are expressed as means ± SEM. *p< 0.05, **p< 0.01, ***p< 0.001 (two-tailed Student’s t-test).

Lack of P2Y_6_R resulted in upregulation of transcript levels coding for *Ppara* and *Pgc1a* in iWAT of adipo-P2Y6^Δ/Δ^ **(Fig. 5C, 5D)**, similar to the effects observed in primary adipocytes lacking P2Y_6_R. Genes involved in mitochondrial function/brown adipocyte function (*Ucp1, Ucp3, Pgc1b, Evovl6, Cox8b, Cytc*, etc.) were upregulated iWAT of adipo-P2Y6^Δ/Δ^ resulting in a switch of white adipocytes to brown adipocyte-like cells **(Fig 5C, 5D)**. Consistent with the gene expression data, protein levels of UCP1, PPARα and PGC1α were dramatically increased in iWAT of adipo-P2Y6^Δ/Δ^ mice **(Fig 5E, 5F)**. In addition, mitochondrial DNA copy number was also significantly increased in iWAT of adipo-P2Y6^Δ/Δ^ mice, consistent with the higher expression of PGC1α, a master regulator of mitochondrial biogenesis **(Fig. 5G)**. In agreement with this observation, the expression of genes coding for mitochondrial transcription factors such as *Nrf1*and *Tfam* and other mitochondrial genes (*Mtco1 and Mtco2*) were also upregulated in iWAT of adipo-P2Y6^Δ/Δ^ mice **(Fig. 5H)**.

Taken together, these experiments demonstrate that white adipocytes lacking P2Y_6_R show reduced JNK signaling resulting in enhanced PPARα-PGC1a activity, upregulation of mitochondrial biogenesis and mitochondria-specific gene expression imparting beige like phenotype to white adipocytes, thereby contributing to reduced inflammation and improved metabolism.

We also carried out transcriptome analysis using total RNA isolated from BAT of adipo-P2Y6^Δ/Δ^ and control mice. The expression levels of genes important for brown adipose function were not significantly different between adipo-P2Y6^Δ/Δ^ and control mice. A list of genes differentially expressed between the groups is given in **Table S2**.

### Generation and metabolic analysis of skeletal muscle-specific P2Y_6_R KO mice (SM-P2Y6^Δ/Δ^ mice)

Having shown that global and adipocyte-specific deletion of P2Y_6_R improves glucose homeostasis, we hypothesized that deletion of P2Y_6_R in skeletal muscle, an important tissue for maintaining glucose homeostasis, may also affect the glucose metabolism. To address this issue, we generated mice lacking P2Y_6_R selectively in skeletal muscle (SM-P2Y6^Δ/Δ^) by crossing P2Y_6_R-floxed mice with mice constitutively expressing the *HSA-Cre* transgene. Expression of HSA-Cre resulted in knockdown of P2Y_6_R mRNA selectively in skeletal muscles **(Fig. 6A)**. Expression of P2Y_6_R mRNA remained unaltered in other major metabolically active tissues **(Fig S5A)**.

**Figure 6:**
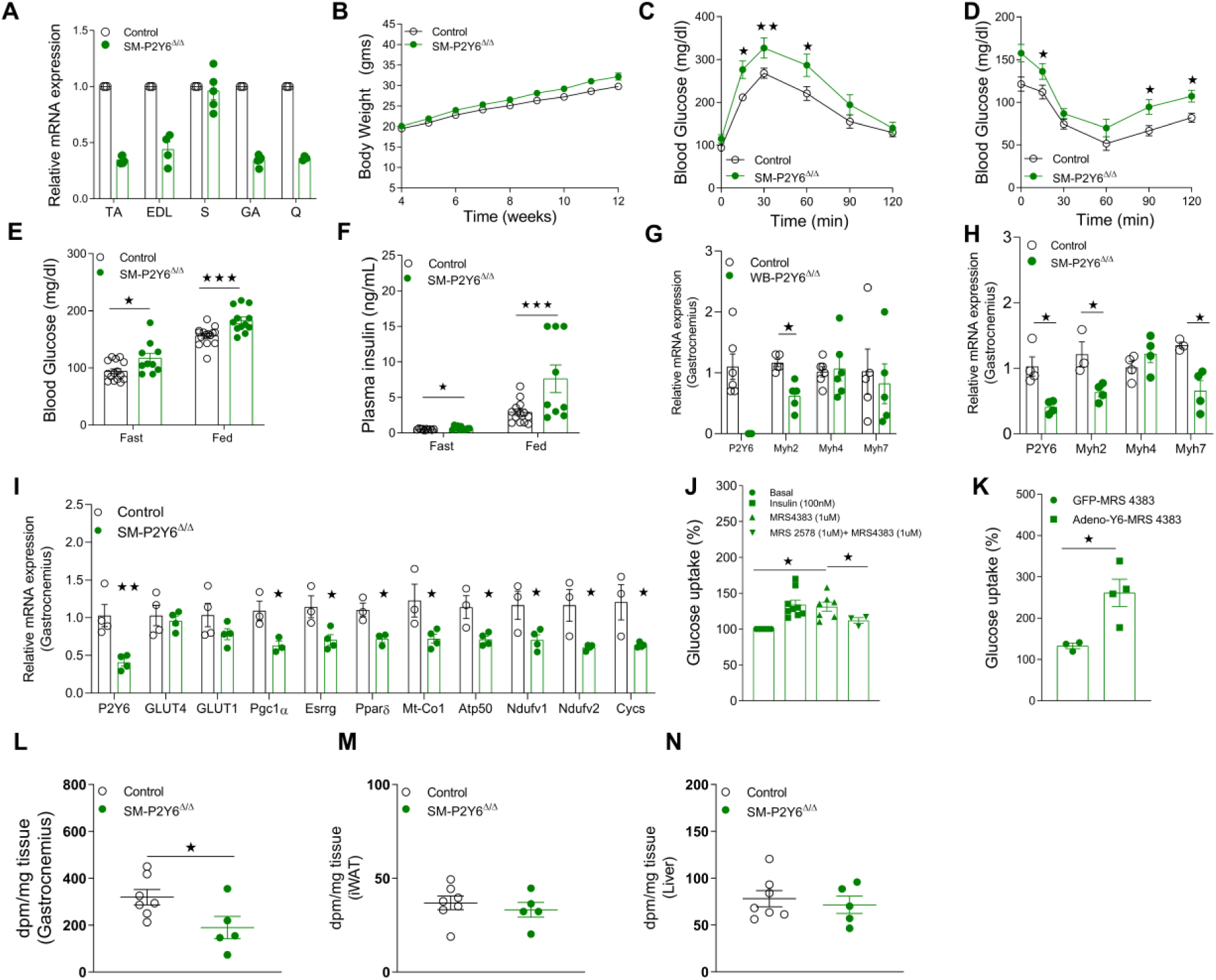
Mice with P2Y_6_R knockout in skeletal muscle (SM-P2Y6^Δ/Δ^ mice) show greater impairments in glucose homeostasis when maintained on HFD. (A) mRNA levels of P2Y_6_R in tibialis anterior (TA), gastrocnemius (G), quadriceps (Q), soleus (S) muscle of SM-P2Y6^Δ/Δ^ and control mice (n=3-4/group). (B) Body weight measurements of mice maintained on HFD (n=9-13/group). (C) Glucose tolerance test (IGTT, 1 g /kg glucose, i.p.) (n=9-13/group). (D) Insulin tolerance test (ITT, 1 U/kg insulin, i.p.) (n=9-13/group). (E) Fasting and fed blood glucose levels (n=9-13/group). (F) Fasting and fed plasma insulin levels (n=9-13/group). (G) mRNA levels of P2Y_6_R and markers of myofiber types in gastrocnemius muscle isolated from WB-P2Y6^Δ/Δ^ and control mice fed with HFD for 16 weeks (n=5-6/group). (H) Quantifying mRNA levels of markers of myofiber types and P2Y_6_R in gastrocnemius muscle isolated from SM-P2Y6^Δ/Δ^ and control mice fed with HFD for 20 weeks (n=3- 4/group). (I) mRNA levels of mitochondrial/oxidative phosphorylation markers in gastrocnemius muscle isolated from SM-P2Y6^Δ/Δ^ and control mice maintained on HFD for 20 weeks (n=3-4/group). (J) Glucose uptake (%) in C2C12 skeletal muscle cells after treatment with insulin (100Nm), P2Y_6_R agonist (MRS4383, 1µM) with or without P2Y_6_R antagonist (MRS2578, 1 µM). (n=3 independent experiments). (K) Glucose uptake (%) in C2C12 skeletal muscle cells transduced with adenovirus expressing GFP or mouse P2Y_6_R. Cells were treated with P2Y_6_R agonist (MRS4383, 1 µM) before quantifying glucose uptake (n=4 independent experiments). (L-N) In vivo ^14^C-2-deoxy-glucose uptake. SM-P2Y6^Δ/Δ^ and control mice fed with HFD for 20 weeks were injected with insulin (Humulin, 0.75 U/kg, i.p.) and a trace amount of [1- ^14^C]2-deoxy-D-glucose (10 μCi). After 40 min, mice were euthanized, and tissues were collected. Quantification of glucose uptake in (L) gastrocnemius, (M) IWAT and (N) liver tissue (n=5-7/group). The expression of 18sRNA was used to normalize qRT-PCR data. All data are expressed as means ± SEM. *p< 0.05, **p< 0.01, ***p< 0.001 (A-B, E-N: two-tailed Student’s t-test; C-D: two-way ANOVA followed by Bonferroni’s post hoc test).

SM-P2Y6^Δ/Δ^ mice and control littermates maintained on chow diet showed no significant differences in body weight, glucose tolerance, insulin sensitivity, blood glucose and plasma insulin levels **(Fig S5B-5F)**. Next, to determine the effect of SM-specific P2Y_6_R deletion on HFD-induced metabolic deficits, we kept mice on HFD starting at 4 weeks of age. SM-P2Y6^Δ/Δ^ mice similar weight compared to control mice on HFD for 8 weeks **(Fig 6B)**. HFD SM-P2Y6^Δ/Δ^ mice displayed significantly impaired glucose tolerance and insulin sensitivity compared to control mice **(Fig. 6C, 6D)**. Consistent with these metabolic deficits, fasting and fed blood glucose as well as plasma insulin levels were increased in SM-P2Y6^Δ/Δ^ mice **(Fig 6E, 6F)**. Somewhat surprisingly, lack of P2Y_6_R in skeletal muscle did not change circulating levels of myokines **(Fig S6A-6E)**. We also studied expression levels of skeletal muscle myofiber markers to examine a potential switch from oxidative to glycolytic myofibers or vice versa. The expression of *Myh4*, a marker for type2b non-oxidative myofibers, was unchanged in skeletal muscle of WB-P2Y6^Δ/Δ^ and SM-P2Y6^Δ/Δ^ mice, as compared to their respective control groups **(Fig. 6G, 6H)**. The mRNA expression levels of *Myh2* and *Myh7*, markers for type 1 and type 2a oxidative myofibers, were decreased in muscle lacking P2Y_6_R **(Fig. 6G, 6H)**. Further, the expression levels of genes related to mitochondrial function/oxidative phosphorylation were decreased in muscle of SM-P2Y6^Δ/Δ^ mice **(Fig. 6I)**. Decreases in oxidative fibers were reported in individuals with obesity or T2D (*32, 33*). Our data suggest that a decrease in oxidative fibers in skeletal muscle of mice lacking P2Y_6_R contributes to impaired glucose homeostasis.

### Activation of P2Y_6_R enhances glucose uptake in skeletal muscle

Since mice lacking P2Y_6_R in skeletal muscle showed significant metabolic deficits, we hypothesized that P2Y_6_R may play a role in glucose uptake in skeletal muscle. To test this, differentiated C2C12 myotubes were stimulated with P2Y_6_R agonist MRS4383 (1 µM). P2Y_6_R activation led to a significant increase in the uptake of radiolabeled 2-deoxyglucose (2-DG). Treatment of cells with P2Y_6_R antagonist MRS2578 (1 µM) prior to stimulation with P2Y_6_R agonist blocked 2-DG uptake by the myotubes **(Fig. 6J)**. We next overexpressed P2Y_6_R in C2C12 myotubes using an adenovirus coding for P2Y_6_R. Cells transduced with a GFP-coding adenovirus were studied for control purposes. Cells overexpressing P2Y_6_R showed enhanced 2- DG uptake compared to the control cells when treated with P2Y_6_R agonist **(Fig. 6K)**. Next, we studied insulin-stimulated *in-vivo* glucose uptake in SM-P2Y6^Δ/Δ^ and control mice maintained on HFD for 20 weeks. As predicted, insulin-mediated glucose uptake in skeletal muscle was significantly lower in SM-P2Y6^Δ/Δ^ mice than in control littermates **(Fig. 6L)**. No difference in insulin-mediated glucose uptake was found in iWAT, BAT, liver and heart of SM- P2Y6^Δ/Δ^ and control mice **(Fig 6M, 6N and S6F-6H)**. These data suggest that activation of P2Y_6_R increases glucose uptake in skeletal muscle independent of the insulin response. In addition, lack of P2Y_6_R *in vivo* renders skeletal muscle resistant to insulin-mediated glucose uptake, thus contributing to impaired glucose homeostasis.

## DISCUSSION

Obesity and T2D epidemic has become a challenging chronic disorder for the countries worldwide. GPCRs have emerged as a major drug targets for various diseases including obesity and T2D (*34*). GPCRs are key players in the control of glucose and energy metabolism, and their function is regulated by various metabolites, neurotransmitters, hormones and nucleotides. Nucleotides and their derivatives activate the P2Y GPCR subfamily. Plasma and tissue levels of these nucleotides fluctuate with energy state and regulate various cell processes through P2YRs and other purinergic receptors (*11*). In the current study, we systematically analyzed the functional role of P2Y_6_R by making global, adipocyte-specific and skeletal muscle-specific KO mice. We demonstrated that loss of P2Y_6_R in mature adipocytes profoundly improved their function in WAT by inducing “beiging” and inhibiting JNK signaling. These changes led to improved systemic glucose metabolism and significantly reduced inflammation. Surprisingly, skeletal muscle P2Y_6_R deletion impaired glucose homeostasis, due to decreased nucleotide-mediated glucose uptake. Moreover, we found that global lack of P2Y_6_R improved glucose homeostasis, suggesting a dominant contribution by adipose tissue P2Y_6_R to the improved systemic glucose metabolism.

P2Y_6_R is expressed in various cell types, and its expression is enhanced by inflammatory stimuli or tissue injury (*35-37*). Feeding mice an obesogenic diet increased P2Y_6_R expression in mature adipocytes, reflecting the receptor’s role in adipocyte-associated pathophysiology in the development of obesity and related metabolic disorders. Here, we show that genetic deletion of P2Y_6_R from mature adipocytes prevented adipocyte hypertrophy and improved glucose tolerance and insulin sensitivity in obese mice. The robust phenotype in adipo-P2Y6^Δ/Δ^ mice was evident in the reduced expression of pro-inflammatory cytokines and lower plasma cytokine levels, suggesting that P2Y_6_R activation causes obesity-associated inflammation in adipose tissue. We also demonstrated that P2Y_6_R deletion resulted in profound molecular changes in WAT. These changes were associated with enhanced “beiging” of WAT with increased expression of genes involved in mitochondrial function/brown adipocyte function (*Ucp1, Ucp3, Pgc1a, Pgc1b, Ppara*, etc). In agreement with the enhanced expression of PGC1α, a master regulator of mitochondrial biogenesis, mitochondrial number was increased in WAT of adipo-P2Y6^Δ/Δ^ mice. This observation supports previous data obtained via DREADD technology, showing that G_q_ activation results in reduced “browning” of WAT (*38*). Another interesting observation was the enhanced expression of sarcoplasmic/endoplasmic reticulum Ca^2+^-ATPase1 (SERCA1-encoded by *Atp2a1*) and ryanodine receptor 1 (*Ryr1*) in WAT of adipo-P2Y6^Δ/Δ^ mice. These proteins are highly expressed in BAT and are involved in calcium cycling between cytosol and the endoplasmic reticulum, releasing energy from ATP hydrolysis in the form of heat (*39*). A previous study reported the role of Serca2 and Ryr2 in UCP1-independent thermogenesis in beige adipose tissue (*40*). Interestingly, no changes in Serca2 expression were observed in our study in WAT of adipo-P2Y6^Δ/Δ^ mice. These findings suggest that lack of P2Y_6_R might lead to Serca isoform-selective calcium cycling in adipocytes. More detailed studies will be required to investigate the possible contribution of P2Y_6_R-mediated regulation of Serca1 expression to thermogenesis in adipocytes. Surprisingly, knockdown of P2Y_6_R in brown adipocytes did not alter the expression of genes involved in brown adipogenesis or thermogenesis. Accordingly, indirect calorimetry did not show any changes in energy expenditure in adipo-P2Y6^Δ/Δ^ mice during the early stage of HFD feeding. The improvement in glucose metabolism observed in adipo-P2Y6^Δ/Δ^ mice may not be detectable during early stages of HFD feeding but may develop after long-term consumption of the HFD, as evident from our study.

Native agonist (UDP)-mediated activation of P2Y_6_R results in IP_3_ accumulation and an increase in cytoplasmic Ca^2+^ levels (*41*). Maintenance of Ca^2+^ homeostasis is essential for the functioning of most metabolic organs including skeletal muscle, pancreas and heart (*42-44*). In adipocytes, alterations in calcium signaling play a major role in the development of metabolic disorder. Rise in cytosolic Ca^2+^ in adipocytes has been shown to promote triglyceride accumulation, reduce lipolysis, inhibit early stages of murine adipocyte differentiation, and promote the production of inflammatory cytokines and immune mediators (*45-47*). In agreement with these studies, mice lacking P2Y_6_R-mediated calcium responses in adipocytes show reduced lipid storage capacity and develop obesity and systemic inflammation. Furthermore, obese individuals show elevated cytosolic Ca^2+^ levels in adipocytes (*48*). In tissues such as liver, increased cytosolic Ca^2+^ levels contribute to JNK activation, inflammatory signaling and enhanced glucose production (*49*). Also, in obesity JNK activity increases significantly in adipose tissue, resulting in severe metabolic impairments (*29*). Mice lacking JNK proteins are resistant to diet-induced obesity, steatosis, inflammation and insulin resistance (*50, 51*). Strikingly, acute P2Y_6_R activation in white adipocytes enhanced JNK phosphorylation, but this effect was absent in adipocytes lacking this receptor, indicating the specificity of JNK activation by P2Y_6_R. Further, P2Y_6_R activation resulted in increased phosphorylation of cJUN which acts downstream of JNK. Interestingly, chronic loss of P2Y_6_R in WAT isolated from adipo-P2Y6^Δ/Δ^ mice resulted in higher protein levels of T-JNK and T-cJUN and increased phosphorylation levels of these proteins. This observation is important, as chronic low-grade inflammation can regulate expression of total JNK in the hypothalamus (*52*).

In addition to JNK activation, we observed that P2Y_6_R stimulation resulted in downstream binding of cJUN to the PPARα promoter, thereby reducing *PPARα* transcription in white adipocytes. Such regulation of *PPARα* transcription by cJUN has also been reported in cardiomyocytes (*53*). We also demonstrated that JNK activation downstream of the P2Y_6_R regulates PPARα transcription, as treatment of cells with JNK inhibitor did not change *Ppara* mRNA levels. However, treatment of cells with JNK inhibitor in the presence of P2Y_6_R agonist rescued the mRNA levels of *Ppara* decreased by receptor activation. P2Y_6_R activation also decreased PPARα activity in white adipocytes, causing decreased expression of PGC1α, a coactivator of PPARα and master regulator of mitochondrial biogenesis **(Fig. S7)**.

We previously reported that P2Y_6_R activation enhanced insulin-independent glucose uptake in C2C12 skeletal muscle cells via AMPK signaling (*15*). In this study, we deciphered the *in vivo* effect of P2Y_6_R deficiency on skeletal muscle glucose metabolism. As expected, mice lacking skeletal muscle P2Y_6_R showed impaired glucose tolerance and insulin sensitivity. In vitro studies confirmed that P2Y_6_R stimulation increased glucose uptake independent of insulin signaling **(Fig. S7)**. These observations are in alignment with a recent study that reported improved glucose metabolism and enhanced glucose uptake in skeletal muscle after activation of a G_q_-coupled designer GPCR (*54*). Furthermore, impaired insulin tolerance in SM-P2Y6^Δ/Δ^ mice was confirmed by decreased insulin-mediated glucose uptake in the gastrocnemius muscle lacking P2Y_6_R.

To understand the interplay and contribution of adipose tissue and skeletal muscle P2Y_6_R on systemic glucose homeostasis in obesity, we studied glucose metabolism in mice with global deletion of P2Y_6_R maintained on an obesogenic diet. Improvement in glucose tolerance and insulin sensitivity were observed with no change in total body weight in WB-P2Y6^Δ/Δ^ mice. These findings are consistent with the outcome of a recent study that reported improved insulin sensitivity in HFD mice lacking P2Y_6_R in whole body or specifically in AgRP neurons of the hypothalamus (*20*). The role of P2Y_6_R in controlling energy balance was also highlighted in a study demonstrating that increased plasma uridine levels in obesity increased hypothalamic UDP synthesis to activate AgRP P2Y_6_R, thereby increasing food intake (*11*). Taken together, these observations indicate that P2Y_6_R activation results in overall negative energy balance and impairment in glucose homeostasis. Importantly, our study provides convincing evidence that global lack of P2Y_6_R improves the metabolic deficits associated with obesity and that adipose tissue is the major contributor to the improved systemic glucose metabolism.

Some limitations of this study should be noted. Due to the short half-life of MRS2578 (P2Y_6_R antagonist) in aqueous medium we could not study the systemic effect over time of selective P2Y_6_R blockade. Furthermore, although we have shown that MRS4383 is a selective agonist of P2Y_6_R, it is not optimally designed for in vivo use, as it is a diphosphate derivative that is subject to hydrolysis by ectonucleotidases. Ongoing efforts in our laboratory are addressing this issue through the synthesis of novel P2Y_6_R ligands. Clearly, this study provides a rational basis for the development of P2Y_6_R antagonists for the treatment of obesity and T2D.

## MATERIALS and METHODS

### Mouse models

Whole body P2Y_6_R KO mice (WB-P2Y6^Δ/Δ^ mice) and mice with a loxP-flanked P2Y_6_R allele (*P2Y6f/f*) have been described (*17*). Mice heterozygous for the P2Y_6_R allele (WB-P2Y6^Δ/WT^) were intermated to generate experimental P2Y_6_R KO mice (WB-P2Y6^Δ/Δ^) and their wild-type (WT) littermates (*WB-P2Y6*^*WT/WT*^, used as control). Mice were maintained on C57BL/6 background.

To generate mice lacking P2Y_6_R selectively in adipocytes, we crossed P2Y_6_R-floxed mice (Dr. Marko Idzko’s lab, Freiburg, Germany) with *adipoq-Cre* mice expressing recombinase under the control of adiponectin promoter. *Adipoq-Cre* mice were purchased from the Jackson Laboratories (Stock No. 010803; genetic background: C57BL/6J). Mice used for experiments were *P2Y6f/f* (control) and *adipoq-Cre*^*+/-*^*P2Y6f/f* (adipo-P2Y6^Δ/Δ^) mice. To generate mice lacking P2Y_6_R selectively in skeletal muscle, we crossed P2Y_6_R-floxed mice with mice expressing recombinase under the control of human skeletal actin promoter (*HSA-Cre 79*; Jackson Laboratories (Stock No. 006149); genetic background: C57BL/6J). Mice used for experiments were *P2Y6f/f* (control) and *HSA-Cre*^*+/-*^*P2Y6f/f* (SM-P2Y6^Δ/Δ^) mice. All mouse experiments were approved by the NIDDK Intramural Research Program Animal Care & Use Committee, Protocol K083-LBC-17.

### Mouse maintenance and diet-induced obesity

Adult male mice were used for all experiments reported in this study. Mice were kept on a 12-h light and 12-h dark cycle. Animals were maintained at room temperature (23°C) on standard chow (7022 NIH-07 diet, 15% kcal fat, energy density 3.1 kcal/g, Envigo, Inc.). Mice had free access to food and water. To induce obesity, groups of mice were switched to a high-fat diet (HFD; F3282, 60% kcal fat, energy density 5.5 kcal/gm, Bio-Serv, Flemington, NJ) at 8 weeks of age. Mice consumed the HFD for at least 8 weeks, unless stated otherwise.

### In vivo metabolic phenotyping

Glucose tolerance tests (IGTT) were carried out in the morning on mice after a 12-h fast overnight. After checking fasted blood glucose levels, animals were i.p. injected with glucose (1 or 2 g/kg as indicated) and blood was collected from tail vein at 15, 30, 60 and 120 min after injection. Blood glucose concentrations were determined using a contour portable glucometer (Bayer). Insulin tolerance test (ITT) and pyruvate tolerance test (PTT) were carried out on mice fasted overnight for 12 h. Fasted blood glucose levels were determined, and then mice were injected i.p. with human insulin (0.75 or 1U/kg; Humulin, Eli Lilly) or sodium pyruvate (2 g/kg) respectively as indicated. Blood glucose levels were determined as described for IGTT. All metabolic tests were performed with adult male mice that were more than 8 weeks of age.

### Body Composition Analysis

Mouse body mass composition (lean versus fat mass) was measured using a 3-in-1 Echo MRI Analyzer (Echo Medical System).

### RNA extraction and gene expression analysis

Tissues from the euthanized mice were dissected and were frozen rapidly on dry ice. Total RNA from tissues under analysis was extracted using the RNeasy mini kit (Qiagen). Total RNA (500 ng of RNA) was converted into cDNA using SuperScript™ III First-Strand Synthesis SuperMix (Invitrogen). Quantitative PCR was performed using SYBR green method (Applied Biosystems). Gene expression was normalized to relative expression of 18s rRNA using the ΔΔCt method. List of primer sequences used in this study are provided in **Supplementary Table S3**.

### Plasma metabolic profiling

Blood was collected from the tail vein of mice in chilled K_2_-EDTA-containing tubes (RAM Scientific). Blood was collected from a group of mice that had free access to food (fed state) or from overnight fasted mice. Collected blood was centrifuged at 4 °C for 10 min at ∼12,000 x g to obtain plasma. ELISA kit (Crystal Chem Inc.) was used to measure plasma insulin levels, following the manufacturer’s instructions. Leptin and adiponectin levels were measured using ELISA kit from R & D Systems. Plasma glycerol, triglyceride, and FFA levels were determined using commercially available kits (Sigma-Aldrich).

### Plasma adipokine/cytokine/myokine levels

Blood was collected from mice fed HFD for 8-10 weeks. Blood was obtained from mandibular vein in a chilled K_2_-EDTA-containing tubes (RAM Scientific). Plasma was obtained by centrifugation of blood at 4 °C for 10 min at ∼12,000 x g. Plasma adipokine and cytokine levels were measured using the Bio-Plex Multiplex Immunoassay System (Bio-Rad), following the manufacturer’s instructions. Plasma myokines were measured using Mouse Myokine Magnetic bead panel (Millipore Sigma). The Luminex Milliplex Analyzer (Luminex) was used to determine the plasma concentrations, as described by the manufacturer.

### Isolation, culture and differentiation of white adipocytes

To isolate mesenchymal stem cells from white fat depot, inguinal white fat pads of 6-8 weeks old male mice was dissected. Excised fat depot was minced into small pieces and digested at 37° C for 45 min in Krebs-Ringer-Hepes-bicarbonate buffer (KRH buffer, 1.2 M NaCl, 40 mM KH_2_PO_4_, 20 mM MgSO_4_, 10 mM CaCl_2_, 100 mM NaHCO_3_, 300 mM HEPES) containing 3.3 mg/ml collagenase I (Sigma-Aldrich). After digestion, 10 ml of KRH buffer was added and cell suspension was filtered through a 70 µm cell strainer, followed by centrifugation at 700 x g for 5 min at room temperature. Supernatant was discarded and the cell pellet was resuspended in KRH buffer and re-centrifuged to obtain cell pellet of mesenchymal stem cells. The pellet was re- suspended in DMEM containing 10% fetal bovine serum (FBS) and 1% pen-strep. Approximately 80,000 cells were seeded per well of collagen I-coated 12-well plates (Corning) and cultured at 37 ° C, 10% CO_2_. Cells were allowed to reach 100% confluency by replenishing media every second day. Two days after reaching confluence, cells were induced for differentiation using DMEM supplemented with 10% FBS, 1% pen-strep, 0.5 µM insulin, 250 µM 3-isobutyl-1-methylxanthine (IBMX), 2 µM troglitazone, 0.5 µM dexamethasone, and 60 µM indomethacin. After 72 h in induction media, cells were incubated in differentiation DMEM media supplemented with 10% FBS, 1% pen-strep, and 0.5 µM insulin for next 72 h. Mature differentiated white adipocytes were used for further studies.

### Isolation, culture and differentiation of brown adipocytes

Brown pre-adipocytes were isolated from interscapular brown BAT of 3-4 weeks old mice. Excised BAT was cut in small pieces and digested at 37° C for 30 min in Krebs-Ringer-Hepes-bicarbonate buffer (KRH buffer, 1.2 M NaCl, 40 mM KH_2_PO_4_, 20 mM MgSO_4_, 10 mM CaCl_2_, 100 mM NaHCO_3_, 300 mM HEPES) containing 3.3 mg/ml collagenase I (Sigma-Aldrich). After digestion, 10 ml of KRH buffer was added and filtered through 70 µm cell strainer, followed by centrifugation at 900 x g for 5 min at room temperature. Cell pellet was washed once in 5 ml of KRH buffer, and cell pellet was re-suspended in DMEM containing 10% fetal bovine serum (FBS) and 1% pen-strep. Cells were seeded in the collagen-I coated 12-well plates (Corning) and cultured at 37 °C, 10% CO_2_. Media was changed every second day until cells reached 100% confluency. Differentiation was induced by added DMEM media supplemented with 2 µg/ml dexamethasone, 0.5 mM IBMX, 0.125 mM indomethacin, 20 nM insulin, and 1 nM T3 to the cells for 48 h. Cells were incubated with DMEM containing 10% FBS, 20 nM insulin, and 1 nM T3 for another 48 h. Mature brown adipocytes were used for experiments.

### Isolation of mature adipocytes

The isolation of mature mouse adipocytes was performed as described previously (*55*). In brief, mouse fat pads were collected and digested with KRH buffer containing collagenase 1 (3 mg/ml). Digested tissues were filtered through a 250 µm cell strainer. After a 5 min centrifugation step at 700 rpm, the top layer containing mature adipocytes was collected and used for further experiments.

### Western blot studies

Adipocyte cells/adipose tissues were homogenized in adipocyte lysis buffer (50 mM Tris, pH 7.4, 500 mM NaCl, 1% NP40, 20% glycerol, 5 mM EDTA, and 1 mM phenylmethylsulfonyl fluoride (PMSF)) supplemented with EDTA-free protease inhibitor cocktail and phosphatase inhibitors cocktail (Roche). Tissue lysates were centrifuged at 14,000 rpm for 20 min at 4°C twice before using the supernatant for western blot studies. Protein concentrations in the lysates were determined using a bicinchoninic acid (BCA) assay kit (Pierce). Protein was denatured in NuPAGE LDS sample buffer (Thermo Fisher Scientific) and β-mercaptoethanol at 90°C for 5 min (*55*). Protein lysates were separated using 4-12% SDS-PAGE (Invitrogen) and transferred to nitrocellulose membranes (Bio-Rad). Membranes were incubated with primary antibody overnight at 4 °C in 5% w/v BSA prepared in 1x TBS with 0.1% Tween 20. Next day, the membranes were washed and incubated with hP-conjugated anti-rabbit/mouse secondary antibody. SuperSignal West Pico Chemiluminescent Substrate (Pierce) was used to visualize the bands on Azure Imager C600 (Azure Biosystems). Images were analyzed using ImageJ (NIH).

### In vivo [^14^C]2-deoxy-glucose uptake

To measure glucose uptake in vivo, mice on HFD for 20 weeks were fasted overnight. Mice were injected with insulin (Humulin, 0.75 U/kg, i.p.) and a trace amount of [1-^14^C]2-deoxy-D-glucose (10 μCi; PerkinElmer). After 40 min, mice were euthanized, and tissues were collected. Tissue weights were recorded, and tissues were homogenized. Radioactivity was measured and counted as described (*56*).

### IP-One quantification

Receptor-mediated intracellular changes in IP1 second messenger were measured using IP-one-Gq kit (Cisbio) as per the manufacturer’s instructions. Briefly, white or brown adipocytes were differentiated in 12-well plates as described before. Cells were serum starved for 4 h and were treated with MRS4383 (1 µM) in stimulation buffer (pH 7.4) containing 50 mM LiCl. After 45 min of stimulation, cells were lysed using 200 µl of lysis buffer (Hepes buffer 1X, 1.5% Triton, O.2% BSA). 14ul of cell lysate was added to the 384-well low volume white microplates (Greiner Bio-One). Working solutions of IP1-d2 followed by Anti-IP1-Cryptate (3 µl) were added to the samples. Plate was sealed and incubated for 1-h at room temperature. Fluorescence was measured at 620 nm and 665 nm with the Mithras LB 940 multimode reader.

### Histology of fat tissues and liver

Different fat depots and liver were dissected out of mice and were quickly frozen in liquid nitrogen. Frozen samples were sectioned and stained using standard techniques. Hematoxylin-eosin (HE) and Oil red O (ORO) stained sections were visualized using Keyence Microscope BZ-9000. Adipocyte images were analyzed for adipocyte size and number using Adiposoft software.

### C2C12 growth and differentiation

C2C12 is a mouse skeletal muscle cell line and is grown and differentiated according to ATCC protocol. Briefly, C2C12 myoblast were cultured at 37° C, 5% CO_2_ in T75 flasks in DMEM supplemented with 5 % FBS, 1% Pen-Strep. Cells were split when after reaching 80% confluency into 12/24 well plates. Media was changed on alternate days until cells reached 80% confluency. Differentiation was started by replacing maintenance media with differentiation media (DMEM supplemented with 2% horse serum, 1% Pen-Strep). Myoblasts were differentiated for 7 days to convert them fully into elongated and contractile myotubes for experiments.

### In vitro glucose uptake in C2C12 cells

C2C12 myotubes were differentiated into myotubes in 12-well plates. 2-Deoxy-glucose uptake in myotubes was measured with modifications as described previously (*15*). Briefly, C2C12 myotubes were serum deprived in DMEM low glucose media for 3 h. Myotubes were treated with P2Y_6_R antagonist (MRS2578, 1,4-di-[(3-isothiocyanatophenyl)-thioureido]butane, 1 µM) in wells as indicated for 30 min. Insulin (100 nM) and P2Y_6_R agonist MRS4383 (1 µM) were added to indicated wells for additional 30 min. After drug treatment, 0.35 ml of transport solution with 0.5 µCi/ml [^3^H]2-deoxy-glucose (PerkinElmer) was added to the cells for 5 min. Cells were washed with ice-cold stop solution (0.9% Saline) and aspirated to dryness. 0.05 N NaOH (1 ml) was added to each well of 12-well plate and radioactivity was measured using scintillation counter. Results were calculated as 2-DG/mg protein/min uptake. Specific uptake in the absence of any stimulation was defined as basal uptake. Data are expressed as percent of basal uptake. To overexpress P2Y_6_R in myotubes, an adenovirus (Ad-GFP-m-P2RY6, Vector Biolabs) was added to myotubes 24 h after inducing differentiation at the MOI of 250. Cells were differentiated for another 5 days before performing glucose uptake studies.

### High throughput RNA sequencing

Adipose tissues were dissected out from mice maintained on HFD for 8 weeks and snap frozen in liquid nitrogen. Total RNA from fat tissues was extracted using the RNeasy mini kit (Qiagen). Quantity and quality of RNA were checked using Agilent Bioanalyzer system. RNA with RIN>8 was used to prepare transcriptomic libraries using the NEBNext Ultra RNA library prep kit (New England Biolabs). High throughput RNA-Sequencing was performed at the NIDDK Genomic Core Facility (NIH, Bethesda, MD) using the HiSeq 2500 instrument (Illumina). Raw sequences were passed through quality control and were mapped to mouse (mm9) genome. Differential gene expression analysis was performed using the Genomatix Genome analyzer with log2 (fold change) cutoff of ±0.58. Biological pathway and enrichment analysis were performed using Metacore (version 6.32, Thomson Reuters, New York) and Ingenuity pathway analysis (Qiagen). The RNA sequencing data have been submitted to the NCBI Sequence Read Archive under the accession code (waiting for the accession number, will be provided later).

### CHIP assay

Differentiated mouse white adipocytes were treated with P2Y_6_R agonist as indicated. After stimulation for 2 h, cells were fixed with formaldehyde, and ChIP assays were performed using the Abcam ChIP kit (ab500) following the manufacturer’s instructions. Antibodies against c-JUN was used to immunoprecipitate cJUN-bound promoter regions. Subsequently, PCR was performed on the eluted DNA using primers annealing to the promoter region of the mouse *PPARa* gene. Primer sequences used for PCR analysis are listed in **Supplementary Table S3**.

### Luciferase assay

Astrocytoma 1321N1 astrocytoma cells stably expressing the human P2Y_6_R were transiently transfected with DR-1, β-galactosidase with or without PPARα, RXRα expressing DNA plasmids using lipofectamine 2000. Cells were treated 48 h after transfection with MRS4383 as indicated. Cells were harvested 6 h after stimulation and luciferase activity was measured according to the manufacturer’s protocol (Promega, Madison, WI). Relative luciferase activity was normalized to the β-galactosidase activity measured using multi-mode plate reader (SpectraMax M5, Molecular Devices) for each sample using manufacturer’s protocol (Promega, Madison, WI).

### Indirect calorimetry

Energy expenditure (O_2_ consumption/CO_2_ production), food intake, respiratory exchange ratio was measured in mice housed at 22°C using an Oxymax-CLAMS (Columbus instruments) monitoring system. Before moving mice to HFD feeding, mice were maintained on CD for three days. On day4, mice were transferred to HFD and data was collected after every 4 min (Total 11 chambers). Mice were maintained on HFD for 4 additional days before ending the experiment. Data Water and food were provided *ad libitum*.

### Statistics

Data are expressed as mean ± s.e.m. for the number of observations indicated. Data were tested by two-way ANOVA, followed by post hoc tests, or by two-tailed unpaired Student’s *t*-test, as appropriate. A P value of < 0.05 was considered statistically significant.

### Study approval

All animal studies were carried out according to the US National Institutes of Health Guidelines for Animal Research and were approved by the NIDDK Institutional Animal Care and Use Committee.

## Acknowledgements

We thank the NIDDK Intramural Research Program for support (ZIA DK311116; ZIA DK311129). We thank Yinyan Ma and Jun Feranil (Mouse Metabolism Core, NIDDK; 1ZICDK070002) for carrying out several metabolite and hormone measurements and indirect calorimetry studies, Dr. Harold Smith (NIDDK Genomics Core) for their help with the RNA-seq work, Dr. Jeffrey Reece (NIDDK Advanced Light Microscopy & Image Analysis Core) provided helpful advice regarding the imaging studies, Dr. Huang Yuning (NIDDK) for tagging and tailing mouse groups.

## Author Contributions

Conceived experimental design: SJ, KAJ; Performed experiments: SJ, SPP, OG; Contributed research materials: KST, BR, MI, JW; Analysis of data and writing of manuscript: SJ, SPP, OG, JW, KAJ; Wrote first draft: SJ.

## Competing interests

The authors declare that they have no competing interests.

## Data and materials availability

All data needed to evaluate the conclusions in the paper are present in the paper and/or the Supplementary Materials. Additional data related to this paper may be requested from the authors.

## REFERENCES

1. D. Segula, Complications of obesity in adults: a short review of the literature. Malawi Med J 26, 20–24 (2014).

2. T. W. Stone, M. McPherson, L. Gail Darlington, Obesity and Cancer: Existing and New Hypotheses for a Causal Connection. EBioMedicine 30, 14–28 (2018).

3. J. O. Hill, H. R. Wyatt, J. C. Peters, Energy balance and obesity. Circulation 126, 126–132 (2012).

4. E. D. Rosen, B. M. Spiegelman, What we talk about when we talk about fat. Cell 156, 20–44 (2014).

5. S. E. Shoelson, L. Herrero, A. Naaz, Obesity, inflammation, and insulin resistance. Gastroenterology 132, 2169–2180 (2007).

6. N. Ouchi, J. L. Parker, J. J. Lugus, K. Walsh, Adipokines in inflammation and metabolic disease. Nat Rev Immunol 11, 85–97 (2011).

7. H. Wu, C. M. Ballantyne, Skeletal muscle inflammation and insulin resistance in obesity. J Clin Invest 127, 43–54 (2017).

8. M. P. Abbracchio et al., International Union of Pharmacology LVIII: update on the P2Y G proteincoupled nucleotide receptors: from molecular mechanisms and pathophysiology to therapy. Pharmacol Rev 58, 281–341 (2006).

9. E. R. Lazarowski, R. C. Boucher, T. K. Harden, Mechanisms of release of nucleotides and integration of their action as P2X- and P2Y-receptor activating molecules. Mol Pharmacol 64, 785–795 (2003).

10. D. Communi, R. Janssens, N. Suarez-Huerta, B. Robaye, J. M. Boeynaems, Advances in signalling by extracellular nucleotides. the role and transduction mechanisms of P2Y receptors. Cell Signal 12, 351–360 (2000).

11. S. M. Steculorum et al., Hypothalamic UDP Increases in Obesity and Promotes Feeding via P2Y6-Dependent Activation of AgRP Neurons. Cell 162, 1404–1417 (2015).

12. P. Badillo et al., High plasma adenosine levels in overweight/obese pregnant women. Purinergic Signal 13, 479–488 (2017).

13. M. A. Laplante, L. Monassier, M. Freund, P. Bousquet, C. Gachet, The purinergic P2Y1 receptor supports leptin secretion in adipose tissue. Endocrinology 151, 2060–2070 (2010).

14. J. Xu et al., GPR105 ablation prevents inflammation and improves insulin sensitivity in mice with diet-induced obesity. J Immunol 189, 1992–1999 (2012).

15. R. Balasubramanian, B. Robaye, J. M. Boeynaems, K. A. Jacobson, Enhancement of glucose uptake in mouse skeletal muscle cells and adipocytes by P2Y6 receptor agonists. PLoS One 9, e116203 (2014).

16. Z. Zhang et al., P2Y(6) agonist uridine 5’-diphosphate promotes host defense against bacterial infection via monocyte chemoattractant protein-1-mediated monocytes/macrophages recruitment. J Immunol 186, 5376–5387 (2011).

17. I. Bar et al., Knockout mice reveal a role for P2Y6 receptor in macrophages, endothelial cells, and vascular smooth muscle cells. Mol Pharmacol 74, 777–784 (2008).

18. A. K. Riegel et al., Selective induction of endothelial P2Y6 nucleotide receptor promotes vascular inflammation. Blood 117, 2548–2555 (2011).

19. M. Nishida et al., P2Y6 receptor-Galpha12/13 signalling in cardiomyocytes triggers pressure overload-induced cardiac fibrosis. EMBO J 27, 3104–3115 (2008).

20. S. M. Steculorum et al., Inhibition of P2Y6 Signaling in AgRP Neurons Reduces Food Intake and Improves Systemic Insulin Sensitivity in Obesity. Cell Rep 18, 1587–1597 (2017).

21. D. Wanders, E. C. Graff, R. L. Judd, Effects of high fat diet on GPR109A and GPR81 gene expression. Biochem Biophys Res Commun 425, 278–283 (2012).

22. S. J. Paulsen et al., Expression of the fatty acid receptor GPR120 in the gut of diet-induced-obese rats and its role in GLP-1 secretion. PLoS One 9, e88227 (2014).

23. D. Osei-Hyiaman et al., Endocannabinoid activation at hepatic CB1 receptors stimulates fatty acid synthesis and contributes to diet-induced obesity. J Clin Invest 115, 1298–1305 (2005).

24. L. Chen, R. Chen, H. Wang, F. Liang, Mechanisms Linking Inflammation to Insulin Resistance. Int J Endocrinol 2015, 508409 (2015).

25. K. S. Toti et al., Pyrimidine Nucleotides Containing a (S)-Methanocarba Ring as P2Y6 Receptor Agonists. Medchemcomm 8, 1897–1908 (2017).

26. J. H. Stern, J. M. Rutkowski, P. E. Scherer, Adiponectin, Leptin, and Fatty Acids in the Maintenance of Metabolic Homeostasis through Adipose Tissue Crosstalk. Cell Metab 23, 770–784 (2016).

27. T. Muller et al., P2Y6 Receptor Activation Promotes Inflammation and Tissue Remodeling in Pulmonary Fibrosis. Front Immunol 8, 1028 (2017).

28. L. Bellocchio et al., Sustained Gq-Protein Signaling Disrupts Striatal Circuits via JNK. J Neurosci 36, 10611–10624 (2016).

29. J. Hirosumi et al., A central role for JNK in obesity and insulin resistance. Nature 420, 333–336 (2002).

30. C. J. Pol et al., Cardiac myocyte KLF5 regulates body weight via alteration of cardiac FGF21. Biochim Biophys Acta Mol Basis Dis 1865, 2125–2137 (2019).

31. D. J. Mangelsdorf et al., The nuclear receptor superfamily: the second decade. Cell 83, 835–839 (1995).

32. J. He, S. Watkins, D. E. Kelley, Skeletal muscle lipid content and oxidative enzyme activity in relation to muscle fiber type in type 2 diabetes and obesity. Diabetes 50, 817–823 (2001).

33. B. Nyholm et al., Evidence of an increased number of type IIb muscle fibers in insulin-resistant first-degree relatives of patients with NIDDM. Diabetes 46, 1822–1828 (1997).

34. F. Reimann, F. M. Gribble, G protein-coupled receptors as new therapeutic targets for type 2 diabetes. Diabetologia 59, 229–233 (2016).

35. S. Koizumi et al., UDP acting at P2Y6 receptors is a mediator of microglial phagocytosis. Nature 446, 1091–1095 (2007).

36. D. M. Grbic, E. Degagne, C. Langlois, A. A. Dupuis, F. P. Gendron, Intestinal inflammation increases the expression of the P2Y6 receptor on epithelial cells and the release of CXC chemokine ligand 8 by UDP. J Immunol 180, 2659–2668 (2008).

37. G. R. Somers, F. M. Hammet, L. Trute, M. C. Southey, D. J. Venter, Expression of the P2Y6 purinergic receptor in human T cells infiltrating inflammatory bowel disease. Lab Invest 78, 1375–1383 (1998).

38. K. Klepac et al., The Gq signalling pathway inhibits brown and beige adipose tissue. Nat Commun 7, 10895 (2016).

39. D. C. da Costa, A. M. Landeira-Fernandez, Thermogenic activity of the Ca2+-ATPase from blue marlin heater organ: regulation by KCl and temperature. Am J Physiol Regul Integr Comp Physiol 297, R1460–1468 (2009).

40. K. Ikeda et al., UCP1-independent signaling involving SERCA2b-mediated calcium cycling regulates beige fat thermogenesis and systemic glucose homeostasis. Nat Med 23, 1454–1465 (2017).

41. A. Brunschweiger, C. E. Muller, P2 receptors activated by uracil nucleotides--an update. Curr Med Chem 13, 289–312 (2006).

42. H. Eshima, D. C. Poole, Y. Kano, In vivo calcium regulation in diabetic skeletal muscle. Cell Calcium 56, 381–389 (2014).

43. A. K. Cardozo et al., Cytokines downregulate the sarcoendoplasmic reticulum pump Ca2+ ATPase 2b and deplete endoplasmic reticulum Ca2+, leading to induction of endoplasmic reticulum stress in pancreatic beta-cells. Diabetes 54, 452–461 (2005).

44. M. Luo, M. E. Anderson, Mechanisms of altered Ca(2)(+) handling in heart failure. Circ Res 113, 690–708 (2013).

45. H. Shi, Y. D. Halvorsen, P. N. Ellis, W. O. Wilkison, M. B. Zemel, Role of intracellular calcium in human adipocyte differentiation. Physiol Genomics 3, 75–82 (2000).

46. M. B. Zemel, H. Shi, B. Greer, D. Dirienzo, P. C. Zemel, Regulation of adiposity by dietary calcium. FASEB J 14, 1132–1138 (2000).

47. I. C. Ho, J. H. Kim, J. W. Rooney, B. M. Spiegelman, L. H. Glimcher, A potential role for the nuclear factor of activated T cells family of transcriptional regulatory proteins in adipogenesis. Proc Natl Acad Sci U S A 95, 15537–15541 (1998).

48. R. L. Byyny, M. LoVerde, S. Lloyd, W. Mitchell, B. Draznin, Cytosolic calcium and insulin resistance in elderly patients with essential hypertension. Am J Hypertens 5, 459–464 (1992).

49. L. Ozcan et al., Calcium signaling through CaMKII regulates hepatic glucose production in fasting and obesity. Cell Metab 15, 739–751 (2012).

50. G. Sabio et al., A stress signaling pathway in adipose tissue regulates hepatic insulin resistance. Science 322, 1539–1543 (2008).

51. G. Tuncman et al., Functional in vivo interactions between JNK1 and JNK2 isoforms in obesity and insulin resistance. Proc Natl Acad Sci U S A 103, 10741–10746 (2006).

52. R. Rorato et al., LPS-Induced Low-Grade Inflammation Increases Hypothalamic JNK Expression and Causes Central Insulin Resistance Irrespective of Body Weight Changes. Int J Mol Sci 18, (2017).

53. K. Drosatos et al., Cardiac Myocyte KLF5 Regulates Ppara Expression and Cardiac Function. Circ Res 118, 241–253 (2016).

54. D. B. J. Bone et al., Skeletal Muscle-Specific Activation of Gq Signaling Maintains Glucose Homeostasis. Diabetes 68, 1341–1352 (2019).

55. S. P. Pydi et al., Adipocyte beta-arrestin-2 is essential for maintaining whole body glucose and energy homeostasis. Nat Commun 10, 2936 (2019).

56. J. K. Kim, O. Gavrilova, Y. Chen, M. L. Reitman, G. I. Shulman, Mechanism of insulin resistance in A-ZIP/F-1 fatless mice. J Biol Chem 275, 8456–8460 (2000).

